# Causal Role of Beta Oscillations in Time Estimation

**DOI:** 10.1101/165233

**Authors:** Martin Wiener, Alomi Parikh, Arielle Krakow, H. Branch Coslett

## Abstract

The neural mechanisms underlying time perception are of vital importance to a comprehensive understanding of behavior and cognition. Recent work has pointed to a supramodal role for beta oscillations in coordinating endogenous timing mechanisms for the purpose of measuring temporal intervals. However, the majority of this work has employed paradigms relying on timed motor responses, which are confounded by beta’s established role in motor movement. Further, no study to date has tested if the alteration of beta oscillations subsequently impacts timing. Here, we address these concerns and demonstrate for the first time a causal connection between beta oscillations and timing. To accomplish this, we first re-analyzed two, separate EEG datasets from psychophysical experiments (Wiener, et al. 2012; 2015) demonstrating that beta oscillations are associated with the retention and comparison of a memory standard for duration, and that transcranial magnetic stimulation (TMS) of the right supramarginal gyrus leads to an increase in midline beta power during the encoding of a temporal interval, corresponding with a longer perceived interval of time. Next, we conducted a study of 20 healthy human participants using transcranial alternating current stimulation (tACS), over frontocentral cortex, at alpha (10Hz) and beta (20Hz) frequencies, during a visual temporal bisection task, demonstrating that beta stimulation exclusively shifts the perception of time such that stimuli are reported as longer in length, while preserving precision. Finally, we decomposed trial-by-trial choice data with a drift diffusion model of decision making and temporal encoding that reveals the shift in timing is caused by a change in the starting point of accumulation, rather than the drift rate or threshold. Our results provide causal evidence of beta’s involvement in the perception of time, and point to a specific role for beta oscillations in the encoding and retention of memory for temporal intervals.

## Introduction

The perception of time stands as one of the primary functions of conscious behavior. Temporal perception underlies a host of cognitive and perceptual functions, and is required for predicting and anticipating upcoming sensory events. However, the neural mechanisms underlying time perception are still being uncovered. To that end, recent research in this area has begun to increase its focus on the role of endogenous oscillations in temporal perception and action (Wiener & Kanai, 2016). Yet, the role of any particular neural oscillation in temporal perception remains to be determined. Indeed, a cross-study survey of neural oscillations in timing reveals that no single frequency band is dominant for temporal processing (Wiener & Kanai, 2016). This finding suggests that distinct frequency bands are context specific to the timing task in which they are employed, but additionally raise the possibility that different frequency bands have different roles in the estimation of a temporal interval. Among these oscillations, recent work has begun to favor a supramodal role for beta oscillations (∼20Hz) in the production and perception of temporal intervals. However, the majority of the evidence in favor of this beta-timing hypothesis is so far circumstantial, and notable challenges to it exist.

The importance of neural oscillations for timing cannot be overstated. Neural oscillations have been proposed as a driving factor for time perception as far back as sixty years (Anliker, 1963), yet failures to replicate these findings led to a reduction of oscillatory studies of timing in the literature, with the majority of studies instead focusing on broadband electrophysiological signatures of timing. Chief among these is the contingent negative variation (CNV), a slow negative potential at frontocentral electrodes that has been associated previously with expectation (Walter, et al. 1964). Studies of the CNV have demonstrated that the amplitude of this response correlates with the subjective perception of temporal duration, suggesting the CNV serves as an index of time for the brain (Macar & Vidal, 2004). Further, the source of the CNV has been suggested as the supplementary motor area (SMA; Casini & Vidal, 2011), a region highly implicated in studies of time perception (Wiener, Turkeltaub, & Coslett, 2010). However, challenges to this interpretation of the CNV exist, driven by a failure to replicate the correlation (van Rijn, et al. 2011) yet, while these failures are troubling, the majority of CNV studies do reveal a close connection between CNV amplitude and subjectively perceived time.

To further investigate the involvement of oscillations in time perception, Kononowicz and van Rijn (2015) re-analyzed an earlier study of the CNV, which had found no association between amplitude and duration (Kononowicz & van Rijn, 2011), by applying a time/frequency analysis of EEG responses. Here, the authors found that the power of beta oscillations exclusively tracked the subjective length of duration. Notably, this task required subjects to produce a temporal interval (2.5s) by indicating via a button press when they believed the interval to have elapsed. The results of this study joined a growing consensus that beta oscillations are intrinsically involved in timing computations. Prior work has demonstrated beta oscillations during temporal preparation and prediction, in which subjects must anticipate the predictable onset of a stimulus (Praamstra, et al. 2006; Fujioka, et al. 2009; Fischer, et al. 2010; Cravo, et al. 2011; Carver, et al. 2012; Arnal, Doelling & Poeppel, 2014). Further, beta oscillations have also been demonstrated during the perception of rhythmic cues, in which beta power and phase track the repeated occurrence of stimuli. Recent neural recordings in non-human primates have also demonstrated beta-oscillations in the basal ganglia associated with motor timing paradigms associated with self-generated rhythmic cues. Notably, beta oscillations exhibit a supramodal property for timing, with similar modulations observed for auditory and visual stimuli.

Yet, challenges to the above interpretation exist. First, all of the above studies entail strong motor components, in which the interval must either be demarcated with a motor response, or the predictable onset of a target stimulus instructs subjects to make a speeded response. As beta oscillations have commonly been associated with motor functions (Baker, 2007), the modulation of beta power in these studies may reflect an upcoming motor response, rather than an actual signal of timing. An exception to the above are studies of rhythmic stimuli (Fujioka, et al. 2009; 2012) in which subjects passively listen to a beat; however, modulation of beta power in these studies may still be driven primarily by cortical and sub-cortical motor areas, as the mere presence of a rhythm is sufficient to activate these areas. Second, the majority of beta studies have employed sub-second temporal intervals. Meijer, te Woerd, and Praamstra (2016) recently demonstrated that, when the anticipatory interval is in the supra-second range, peaks in beta resynchronization occur at the same point in time, between 800-1000ms, regardless of how long the anticipatory interval lasts. Yet, this study primarily demonstrates a potential dissociation between motor preparatory processes and cannot rule out entirely that differences observed between interval duration conditions do not reflect timing computations. Further, the study by Kononowicz and van Rijn (2015) included supra-second intervals (∼2.5s), suggesting the difference may have relied on the implicit/explicit nature of the timing task (Coull & Nobre, 2008).

Only one study that we are aware of has addressed the beta timing hypothesis in a non-motor paradigm. Kulashekhar and colleagues (2016) tested subjects with MEG on a temporal discrimination task with supra-second durations, using the well-known time/color paradigm of Coull and colleagues (Coull, et al. 2004), in which subjects view a visual stimulus rapidly flickering between shades of red and blue, and must either compare the duration or overall color of the stimulus. Here, the authors demonstrated that duration comparisons engendered significantly greater beta power amplitudes in a parieto-frontal network – including the SMA – during the encoding of stimulus duration. Further, higher beta amplitudes were associated with accurate vs. inaccurate choices, suggesting strong behavioral relevance.

Beyond beta oscillations, as noted above, other frequency bands have been associated with timing functions (Wiener & Kanai, 2016). The second most associated frequency band to beta oscillations are slower, alpha oscillations. Indeed, alpha oscillations were the first sort to be associated with time perception (Anliker, 1963), yet their association could historically not be replicated (Legg, 1968). More recently, alpha oscillations have been observed in visual temporal prediction studies (Praamstra, et al. 2006; Rohenkohl & Nobre, 2011; Samaha, et al. 2015). For non-motor timing paradigms, alpha oscillations have also been observed (Ng & Penney, 2011; Kulashekhar, et al. 2016). In the study by Kulashekhar and colleagues, alpha oscillations were stronger during the retrieval of duration information from memory, and were similarly associated with accurate over inaccurate decisions, suggesting that alpha power in this task may reflect decision making for temporal duration. Further, recent work has suggested that alpha frequency serves as an index of the “sample rate” for the visual system, such that fluctuations in peak alpha frequency may be associated with changes in perceived duration (Samaha & Postle, 2015; Cecere, Rees, & Romei, 2015).

While the highlights above demonstrate the association of beta and alpha oscillations with time perception and temporal prediction, none of the aforementioned studies have demonstrated that beta oscillations are causally associated with timing. That is, the involvement of either of these frequency bands may be epiphenomenal and not reflect actual timing computations. For this, interventional methods are necessary to perturb a particular frequency band and observe if temporal perception is commensurately affected. Furthermore, the particular way in which timing may be changed as a result of this perturbation can be informative the precise mechanism that has been affected. Time perception may be fractionated into distinct cognitive or perceptual states, such that the perception of time must entail encoding, storage, retrieval and comparison processes (Gibbon, Church & Meck, 1984; Matell & Meck, 2000); different oscillatory frequencies may similarly be associated with these different stages (Kulashekhar, et al. 2016).

To address the above issues and directly test the beta-timing hypothesis, we tested subjects on a sub-second visual temporal bisection task, in which subjects must categorize stimulus durations into “long” and “short” categories (Wiener, Thompson & Coslett, 2014). To directly test the causal contribution of beta oscillations, we applied transcranial alternating current stimulation (tACS), a type of noninvasive electrical stimulation in which the frequency can be specified, at both beta (20Hz) and alpha (10Hz) frequencies in separate sessions. Studies of tACS have demonstrated the efficacy of 1 milliamp stimulation at entraining neural populations to a specified frequency and increasing the power at that band. Further, as few studies have investigated the role of beta oscillations during non-motor time perception tasks, we re-analyzed two previously collected EEG datasets (Wiener, et al. 2012; Wiener & Thompson, 2015) recorded from tasks similar to the present one. In one (Wiener & Thompson, 2015), subjects were tested on an auditory sub-second temporal bisection task of the exact same design as the one used in the present study. This study was originally an investigation of the CNV signal and its role in memory for temporal intervals, demonstrating that the amplitude of the CNV signal on a trial-by-trial basis reflected both the duration of the present trial and changes in the average perceived duration across trials. In the other (Wiener, et al. 2012), subjects performed a visual sub-second temporal discrimination task, while repetitive transcranial magnetic stimulation (TMS) was applied to the right supramarginal gyrus (SMG) of the parietal cortex, or to a control site in the midline at the junction of the occipital and parietal cortices, prior to the onset of the standard stimulus (600ms). This study also focused on the CNV, demonstrating that stimulation of the SMG led to an increase in amplitude during the presentation of the standard stimulus, but with no change during the comparison stimulus. Further, the increase in CNV amplitude paralleled a shift in perception, such that subjects perceived the standard stimulus as longer in duration, with the size of both effects correlating between subjects. Accordingly, in re-analyzing these two datasets, we hoped to identify whether beta oscillations were implicated in time perception, as previously demonstrated by other studies. Further, the re-analysis allowed us to investigate timing in two non-motor paradigms, such that the report of stimulus duration was not dependent on the timing of the motor response.

Lastly, in order to identify precisely which aspect of time perception may have been influenced by the application of tACS, we decomposed choice and reaction time data from the temporal bisection task with a drift diffusion model (DDM) of decision making (Ratcliff, 1978; Wiecki, et al. 2013). Recent work has demonstrated that temporal perception may be constructed as a variant of the DDM, known as the Time-Adaptive Opponent Process Drift Diffusion Model (TopDDM; Simen, et al. 2011), in which opponent populations of neurons provide a bound for temporal accumulation. An extension of this model to decision making in 2AFC designs (Balci & Simen, 2014) has demonstrated that the standard DDM, in which decision evidence accumulates to one of two boundaries at a particular rate, can well describe performance on the temporal bisection task. As such, particular elements of the DDM can be tied to distinct processing stages in time perception. We here applied a hierarchical DDM (HDDM; Wiecki, et al. 2013), in which Bayesian posterior estimates of model parameters are determined via Markov Chain Monte Carlo (MCMC) sampling, and the individual subject estimates are constrained, hierarchically, by the mean group estimate. This procedure has been successful previously at describing performance on temporal bisection (Tipples, 2015) with findings predicted by TopDDM (Balci & Simen, 2014).

## Methods

### Subjects

20 Healthy, right-handed subjects (12 male; Mean age: 24 ± 4 years) participated in the tACS experiment. One subject was removed due to an error in data collection where the same stimulation frequency was administered twice, resulting in an *n* of 19. All subjects were tested at the University of Pennsylvania, and drawn from the general population of the area via flyers. Exclusion criteria were 1) history of active neurological or psychiatric disease, 2) history of seizures or unexplained consciousness, 3) history of brain surgery, craniotomy, or other breach of the skull, 4) current consumption of anti-convulsant, anti-psychotic, or sedative/hypnotic medications, 5) participation in other brain stimulation experiments on the same day, 6) prior adverse reaction to non-invasive brain stimulation, 6), metallic hardware in or on the head that could not be removed (excluding dental work), and 7) pregnancy. All subjects gave their informed consent as approved by the University of Pennsylvania Institutional Review Board. A neurologist was on-call during each session.

### Task Design

All subjects performed a visual temporal bisection task, using the ‘partition method’ (Wearden & Ferrara, 1995). Task design was identical to our earlier reports using this task (Wiener, et al. 2014; Wiener & Thompson, 2015). In the temporal bisection task, subjects must classify a stimulus, presented for a specific duration, into either “long” or “short” duration categories (Figure 1a). On a given trial, following a 500ms fixation point, subjects were presented with a Gaussian luminance blur, presented at 100% contrast against a grey background with a FWHM of 2cm. The visual stimulus could persist for one of seven logarithmically spaced durations between 300 and 900ms. Following each presentation, subjects were required to classify the stimulus with a button press. Subjects were specifically instructed to classify each stimulus duration by comparing it to the average of all previously presented stimuli. To assist in the classification for the first few trials, each session began with three stimulus presentations at the geometric mean of the stimulus set (520ms). Following the offset of the stimulus, subjects were instructed to provide a response as quickly, but as accurately, as possible. Similar to our previous reports, the order of stimulus presentation was determined by a path-guided de Bruijn sequence (Aguirre, Mattar, & Magis-Weinberg, 2011). Briefly, the de Bruijn sequence allowed us to present all stimulus durations in a sequence that was first-order counterbalanced, such that every possible transition from one duration to another occurred an equal number of times. This allowed us to accurately measure the influence of carryover effects from the previous trial. An additional label for null (empty) trials was added to the sequence, so as to include trial transitions where no stimulus was presented; on null trials, subjects viewed a blank screen for 550ms, followed by the appearance of the fixation point for the next trial. The resulting trial matrix consisted of 64 possible trial types and a sequence of 448 trials (excluding nulls). Each duration in the total sequence was presented 64 times. The time to run through a single sequence was ∼15 minutes.

**Figure 1.**
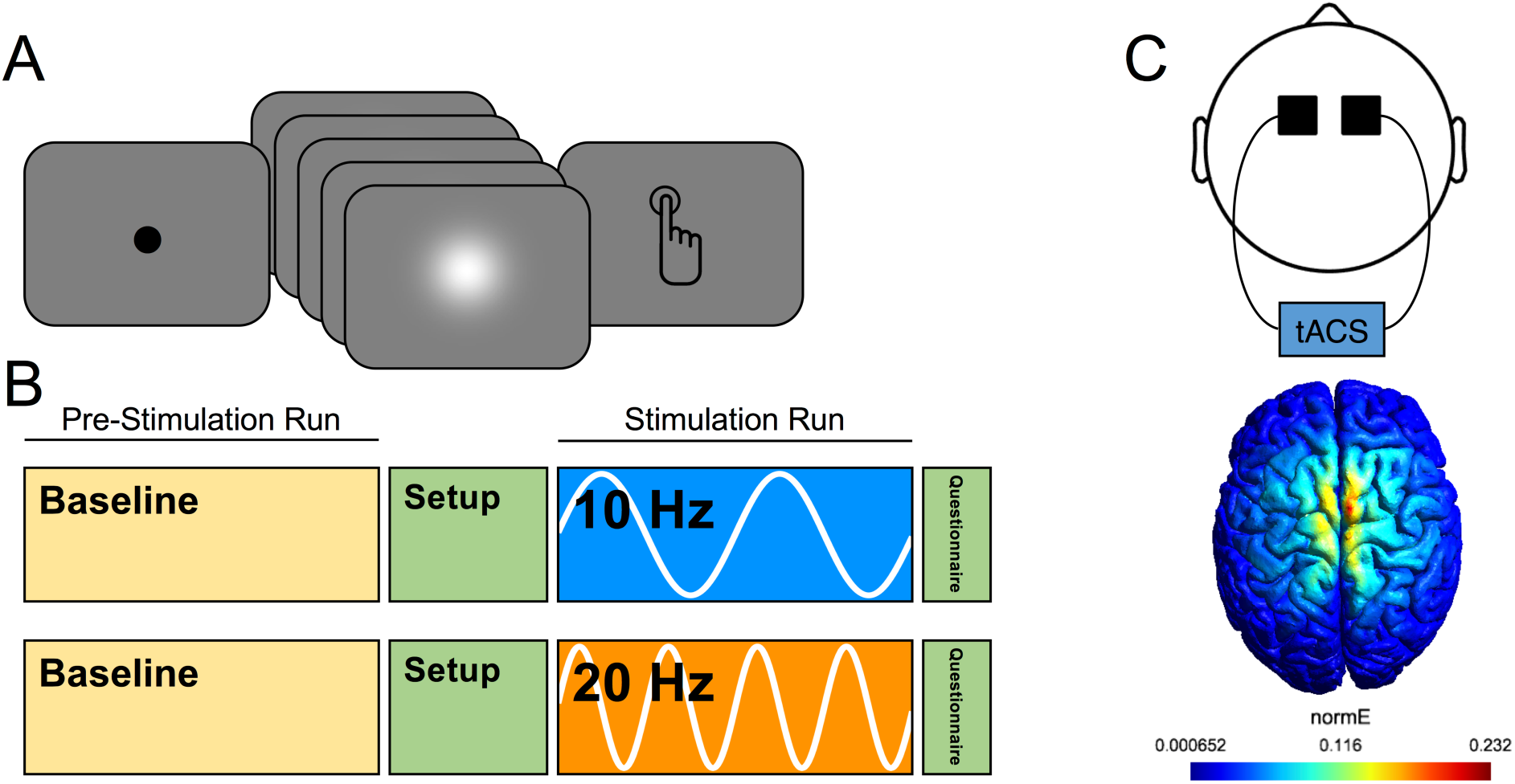
**A**) Task design for visual temporal bisection. Subjects viewed a fixation point for 500ms, and where then presented with a visual stimulus (Gaussian luminance blur) for one of seven possible durations, logarithmically-spaced between 300 and 900ms. At offset, subjects were required to classify the presented stimulus into “long” or “short” duration categories. **B**) Experimental setup. All subjects (N=20) started each session by performing a baseline version of the temporal bisection task. Following this, tACS electrodes were administered on the subject and stimulation was initiated; all subjects performed a baseline questionnaire at this time. In the stimulation run, subjects again performed the bisection task, while concurrently receiving stimulation (10Hz or 20Hz, separate days). Following stimulation, subjects again filled out a questionnaire. **C**) At top, the electrode montage used for tACS, with electrodes placed over FC1 and FC2 in the international 10-20 system. At bottom, the results of electric field modeling for our chosen montage, demonstrating a maximal electric field generation within the supplementary motor area.

### tACS

The study was a 2-session single-blind experiment, with 2 days between sessions and verum stimulation at either 10 Hz or 20 Hz applied during each session (Figure 1b). At each session, participants first completed the bisection task with no stimulation. Positions of FC1 and FC2 were then determined using the 10-20 system. These electrodes were chosen to maximize bilateral stimulation of the SMA (see simulation details below). 5 × 5 cm rubber electrodes placed inside sponges soaked in a 0.9% sterile saline solution (MOLTOX) were placed over FC1 and FC2 and held in place with a rubber headband. Impedance was reduced if the safety criteria of the tACS device (Neuroconn Magstim DC-Stimulator) were not met by soaking hair in saline and tightening the headband to improve electrode contact with the scalp. tACS was delivered at 1.5 mA peak to peak and at either 10 Hz or 20 Hz. Participants completed the bisection task once again while they were receiving stimulation, beginning five minutes after the initiation of stimulation. Participants were stimulated for a total of 20 minutes. The tACS frequency of the first session was pseudo-randomized and counterbalanced.

Subjects completed a post-study questionnaire after each session to assess side effects (tingling, itching sensation, burning sensation, pain, fatigue, nervousness, headache, difficulty concentrating, mood change, change in visual perception, visual sensation, or other effects) during and after tACS, using a visual analog scale to rate each category from 0 to 10. After the second session, they were also asked whether they believed they were receiving verum or sham stimulation during each session.

### Simulation

To determine the likely physiological source of stimulation, we simulated the effect of our electrode montage using the SimNIBS 2.0 Toolbox (http://simnibs.de/; Thielscher, et al. 2015). As in our main experiment, the placement of two electrodes (5 × 5cm) located at FC1 and FC2, were simulated with a current of 1.5mA. The normalized electrical field was simulated via a realistic finite element head model (Opitz, et al. 2013). As both frequencies used the same stimulation amplitude, only one simulation was run (Moisa, et al. 2016).

### Behavioral Analysis

Behavioral analysis of the temporal bisection task follows earlier reports (Wiener, et al. 2014; Wiener & Thompson, 2015). Choice and reaction time data were first calculated for each subject for each of the seven tested durations in our stimulus set. Trials were then filtered to only include those where the RT was between 100 and 1000ms, to eliminate premature and late responses (see supplementary figure 1 for average # trials removed). The justification for this cutoff was to ensure our ability to measure carryover effects from the previous trial (Wiener, et al. 2014). Psychometric and Chronometric curves were then generated for each participant. Psychometric curves were generated by plotting the proportion of long response choices for each of the seven tested durations; these points were then fitted by a cumulative Gumbel distribution using the *psignifit* version 3.0 software package (see http://psignifit.sourceforge.net/) for Python (Fründ, Haenel, & Wichmann, 2011). The Gumbel distribution was chosen to reflect the log-spacing of the duration stimuli, as well as the uncertainty associated with longer stimulus durations (van Driel, et al. 2014). Upper and lower thresholds, the approximate points at which the subject is 25% or 75% likely to judge the stimulus as long, were calculated using the bias-corrected bootstrap method implemented by psignifit, based on 1999 simulations. The results of this analysis yielded the bisection point (BP; the time value at which subjects were equally likely to judge the stimulus as long or short), the difference limen (DL; the difference between the upper [75%] and lower [25%] threshold values divided in half), and the coefficient of variation (CV; DL/BP). The BP thus reflects the subjective midpoint of the range of tested durations, while the CV reflects the normalized variability of measurements. Chronometric curves were constructed by plotting the RT for each of the seven possible durations.

The use of a de Bruijn sequence in our study, as in our prior work (Wiener et al. 2014; Wiener & Thompson, 2015) allows us to investigate the potential impact of carryover effects. Briefly, psychometric and chronometric curves were generated as described above for each of the seven possible *prior* durations that could have occurred on the previous trial. A positive carryover effect is characterized by a contrast of the present duration away from the duration presented on the previous trial. As such, the BP is expected to shift leftward for trials on which the previous trial was long, and rightward for trials on which the previous trial was short. This can be characterized by the slope of a linear regression of BP values across all seven prior durations. The change in the BP is expected to reflect fluctuations in the mean remembered duration, or criterion value for classifying durations into long and short categories. Additionally, bisection choice data also exhibits a gravitation of the current decision towards the decision made on the previous trial. As such, the BP is shifted to the left if the prior trial’s response was “long” and to the right if the previous trial’s response was “short”, regardless of the duration presented on that trial. This change can be characterized by the difference in BP values between data sorted on the basis of the prior trial response. Both effects were investigated in the present study.

### Drift Diffusion Modeling

In order to further parse the source of tACS effects on behavior, choice and reaction time data were decomposed with a drift diffusion model (DDM). In this case, we sought to implement the TopDDM, as described by Balci and Simen (2014). In the original formulation of the TopDDM (Simen, et al. 2011), the model assumes that populations of neurons with Poisson firing rates serve as Clock Pulse Generators. Spikes from these generators are integrated by readout neurons over time, while simultaneously inhibited by interneurons that are also triggered by the generators. This process serves to allow for linear accumulation in the integrator neurons, and can be mathematically approximated as a stochastic differential equation *dx* = *A* ∙ *dt* + m ∙ 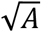 ∙ *dB*. The TopDDM thus assumes that, during a timed interval, accumulator neurons integrate the opponent processes, which serves as an elapsed measurement of time for a particular accumulation rate (*A*). This measurement elapses until the accumulated output crosses a fixed threshold. Notably, TopDDM assumes that, once subjects learn the duration to be timed, the accumulation rate *adapts*, via a learning rule, such that the drift is higher for shorter durations and lower for longer durations. In the extension to temporal bisection (Balci & Simen, 2014), the model assumes that once the presented stimulus extinguishes, a second-stage drift diffusion process (*v*) is triggered that accumulates towards one of two decision boundaries (*a*). In an elegant extension, the starting point and direction of this second, decision drift process is determined by the end point of the first, accumulator drift process (Figure 2). Application of the two-stage TopDDM process to temporal bisection data thus provides a decomposition of both accumulator and decision processes during timing.

**Figure 2.**
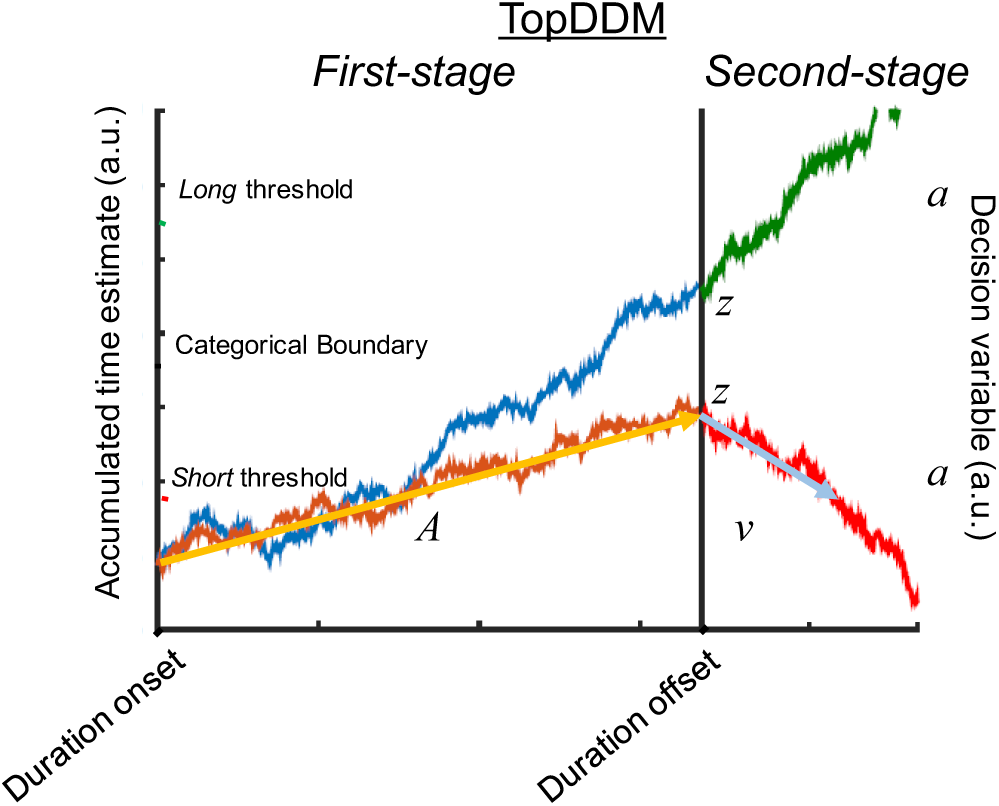
Example of the TopDDM model for temporal bisection (cf. Balci & Simen, 2014). On a given trial, at the onset of a to-be-timed interval, a first-stage drift process is initiated that accumulates at a particular rate (A) until duration offset occurs. At offset, a second-stage is initiated, with the ending accumulated time estimate of the first-stage serving as the starting point (z). From here, a decision variable accumulates towards one of two decision boundaries (a) at a particular rate (v). The direction of the drift depends on where the starting point lies relative to a categorical boundary that is used for classifying stimuli. In the example above, the same interval may be categorized as short or long, depending on the first-stage drift rate.

In the present study, we focused our modeling effort solely on the second-stage process. Justification for this comes from results demonstrating that modeling the second-stage process alone is sufficient to account for bisection data and exhibits the same properties as if both stages had been modeled (Balci & Simen, 2014; Tipples, 2015). These properties include: 1) the drift rate (*v*) should be negative for durations shorter than the BP and positive for durations longer than the BP; 2) The non-decision time (*t*) should decrease linearly for longer durations; 3) the starting point (*z*) should linearly increase for longer durations; 4) the variability of the starting point (*sz*) should increase for longer durations; 5) The threshold (*a*) should decrease for durations closer to the BP. To accomplish this, we applied hierarchical Bayesian estimation for the by using routines provided by the HDDM toolbox for Python (Wiecki, et al. 2013), which implements a Bayesian inference, via Markov chain Monte Carlo (MCMC) sampling, for inferring and selecting model parameters. HDDM estimates individual subject parameters by accounting for group-level response distributions (Wiecki, et al., 2013). Markov Chain Monte Carlo (MCMC) methods were then used to determine posterior probability distributions for each parameter included in the model. As described above, we decomposed choice and reaction time data to obtain four measures: threshold (*a*), drift-rate (*v*), non-decision-time (*t*), and starting point (*z*).

To account for model convergence, several steps were taken. First, model comparisons were conducted between models of varying complexity. We sought to compare a factorial expansion of our experimental design, and so we compared performance of an “empty” model, in which all model parameters varied only by subject, to three models of increasing complexity (frequency, frequency × stimulation, frequency × stimulation × duration). Model comparison was conducted using the deviance information criterion (DIC; Spiegelhalter, et al. 2002); a DIC difference greater than 10 was taken as evidence in favor of a more complex model. Each model was estimated using a chain of 10,000 samples from the posterior distribution, with the first 1000 samples discarded as burn-in, and only every 5^th^ sample thereafter retained; this process, known as “thinning”, is used to increase chain stability and reduce autocorrelations between subsequent samples. Once the “winning” model was found, we further sampled five additional chains of 5000 iterations (200 burn-in) and compared the Gelman-Rubin Statistic (Gelman & Rubin, 1992). Finally, we conducted posterior predictive checks (PPC; Kruschke, 2013) by using model parameters to simulate data from 500 observers and compare those data to our original set of subjects.

One aspect of the modeling approach here is that the parameters for each subject, in each condition, are estimated hierarchically, such that the group distribution is used to constrain values for individual subject and trial-wise parameters. We therefore sought to determine if this approach influenced our results. To do this, we also estimated DDM parameters in a “non-hierarchical” fashion, by applying our HDDM model to each subject individually, and then combining model values. We again compared factorial model complexity, to ensure that the winning, hierarchical model was not driven by the method of fitting. In this case, we compared the average Akaike Information Criterion (AIC) score between models, as the DIC was developed the assess hierarchical models only; further, AIC models can be compared by measuring the relative likelihood of each model compared to the model with the lowest AIC value (Burnham & Anderson, 2002).

Lastly, we also explored the application of a Hierarchical Generalized Linear Model (GLM) approach to HDDM, by applying the HDDMRegressor() class of the HDDM toolbox. Here, the involvement of model parameters can be directly regressed onto choice and reaction time data. This approach allows for the direct comparison, via Bayesian statistics, of different posterior distributions. Again, model complexity was explored to pick the winning model (via DIC). In this case, the frequency of stimulation and stimulation status (baseline vs stim) were modeled as dummy variables, with [alpha,baseline] as the intercept in the most complex model. Duration was modeled as a continuous covariate. Parameters for *v, t,* and *a* were modeled as linear relations, whereas *z* was modeled by an inverse logit link function (see http://ski.clps.brown.edu/hddm_docs/tutorial_regression_stimcoding.html#chap-tutorial-hddm-regression for details on this process) to constrain parameter values between 0 and 1. Comparisons were done on the group posterior distributions to assess significance (Krushcke, 2013).

### EEG Re-analysis

*Wiener & Thompson (2015)*. In order to establish what role, if any, beta oscillations may play in temporal perception, we first re-analyzed data from a previous temporal bisection EEG study (Wiener & Thompson, 2015). This study consisted of an auditory temporal bisection task, using the exact same durations and setup as in the present study, with the exception that an auditory stimulus (white noise burst) was used instead of a visual disc. Subjects performed three runs, consisting of 512 trials each. EEG data were collected via 64-channel setup (Synamps^2^) at 500Hz with a bandpass filter of 0.1-100Hz. All preprocessing steps were identical to our earlier study for these data; briefly, the data were rereferenced to the average of two Mastoid electrodes, noisy channels were removed and interpolated, and a trialwise cutoff of ±120μV was used to eliminate trials with excessive noise. Data were further epoched into both onset-locked (−400 – 1000ms) and response-locked (−1000 – 200ms) data.

To examine time/frequency effects, we applied a Morlet wavelet convolution, via the EEGLAB *newtimef* function, by convolving a mother wavelet at 100 log-spaced frequencies spanning 10 to 40 Hz, with 3 cycle wavelets and a 0.5 scaling factor. To examine activity immediately prior to the response, we calculated the difference between spectral power for trials on which the subject classified the stimulus as “long” vs “short”. Significance was assessed via a cluster-based permutation statistic (Maris & Oostenveld, 2007), based on the maximum cluster size of paired-samples *t*-tests at each time/frequency point. This procedure most adequately controls for the false-positive rate in cluster inference (Pernet, et al. 2015). For onset-locked data, we separately examined the impact of the present trial duration (termed *direct effect*) and the prior trial duration, collapsing across all present trial durations (termed *carryover effect*). Our original study had demonstrated that the frontocentral CNV signal, maximal at FCz, covaried linearly with both the present and prior trial durations. As such, we focused our analysis on FCz. To measure direct and carryover effects, we modeled the impact of duration as a linear regression at each time/frequency point with duration. The resulting slope values of the best fitting regression line indicate if power values at a particular time/frequency linearly increased or decreased with duration. These slope values were then tested for significance with a one-sample *t*-test versus zero (no change with duration) with a cluster correction for multiple comparisons, as described above.

*Wiener, et al. (2012).* As a second means of establishing what role, if any, beta oscillations play in temporal perception, we also re-analyzed data from a study by Wiener and colleagues (2012). In this study, nineteen participants performed a visual temporal discrimination task, wherein a comparison duration must be judged as longer or shorter than a standard duration (600ms) presented on each trial. The visual stimulus used was a 4x4cm red square; the comparison durations were individually titrated via a baseline adaptive staircase procedure, and included three possible values: 600ms and the upper and lower thresholds for determining a stimulus as different than 600ms. EEG was collected via a 32-electrode high-impedance system (BioSemi) at 512Hz. Additionally, a brief train of repetitive TMS was administered at 10Hz (3 pulses), at 500ms prior to the onset of the standard duration. A single run consisted of 180 trials, half of which randomly included TMS. Two separate sessions on separate days were conducted, in which TMS was administered to either the right supramarginal gyrus (rSMG) or the midline-occipital parietal cortex (Mid-Occ). Target locations were individually determined via high-resolution T1-weighted anatomical images and localized using Brainsight (Rogue Research). The rSMG target was approximately over electrode CP6, whereas the Mid-Occ target was approximately over electrode Pz. Each of these electrodes were unplugged and removed during the stimulation session to reduce the distance between the TMS coil and the scalp. Each electrode was interpolated in the resulting EEG analysis.

Preprocessing steps were similar to the previous study, with some exceptions. Data were again offline referenced to the common average of all available channels, with noisy channels removed and interpolated, but a wider offline bandpass filter (infinite impulse response) was applied between 1 and 50Hz. Data were epoched between −700 and 1000ms to the onset of the standard stimulus with a ±90μV cutoff to eliminate noisy trials. This wider epoch, necessary to adequately resolve time/frequency effects for the baseline, included the artifacts induced by TMS. To resolve this, we used a similar strategy for reducing and eliminating the TMS artifact as our previous study by the application of a spatial filter with independent component analysis (ICA). The steps for artifact removal are similar to those outlined by Herring and colleagues (2015). Specifically, the space of EEG spanning the artifact (−500 to −200ms prior to standard stimulus onset) was removed from each trial. Infomax ICA was then applied to the data in order to identify components related to residual artifacts resulting from muscle artifacts and the return to baseline. ICA components related to resulting artifacts as well as eye-blinks were determined on a per subject/session basis by visual inspection of the topography and power spectrum of the component. Noise components were then used as a spatial filter for the data by their removal and projection back to the original dataset. Finally, the lost space between −500 and −200ms was interpolated. Visual inspection of our data confirmed that the ERP results looked qualitatively similar to our original report, with a larger CNV amplitude for rSMG stimulation compared to Mid-Occ stimulation at electrode Cz.

For the analysis of time/frequency effects, we focused our analysis on the stimulation condition only for each condition. Again, we applied a wavelet approach, using similar parameters to the above re-analysis. One difference for this analysis from above is that we applied a trialwise baseline (−200 to onset) via single-trial division. We opted for this method of baseline correction, instead of the standard method of averaging spectral estimates before baseline removal, as it has been shown to reduce the sensitivity to noisy trials in time/frequency analyses (Grandchamp & Delorme, 2011). As the number of trials in each condition was low (*n=*90), we wanted to avoid contamination by noise or outliers. Significance was assessed by calculating the difference between rSMG and Mid-Occ conditions at electrode Cz via a paired-samples *t*-test, cluster corrected.

## Results

### EEG Re-analyses

The majority of research demonstrating an effect of beta frequency power on temporal processing have utilized motor-based timing paradigms, wherein the motor response determines the interval being measured or entrained. To begin examining the impact of distinct frequency bands in timing, we first re-analyzed a prior dataset (Wiener & Thompson, 2015) using an auditory temporal bisection task with sub-second stimuli. In this study, a difference in the amplitude of a frontocentral CNV signal was observed in response-locked data, depending on whether a subject classified a stimulus as “long” or “short”. The spectral decomposition of these data revealed a significant event-related desynchronization (ERD) occurring 500ms prior to the response, followed by a significant event-related synchronization (ERS) centered on the response, for trials on which subjects judged the stimulus as long, compared to short (Figure 3a). The frequency space of these changes in power spanned across alpha and beta frequency ranges, but was larger in the beta range.

**Figure 3.**
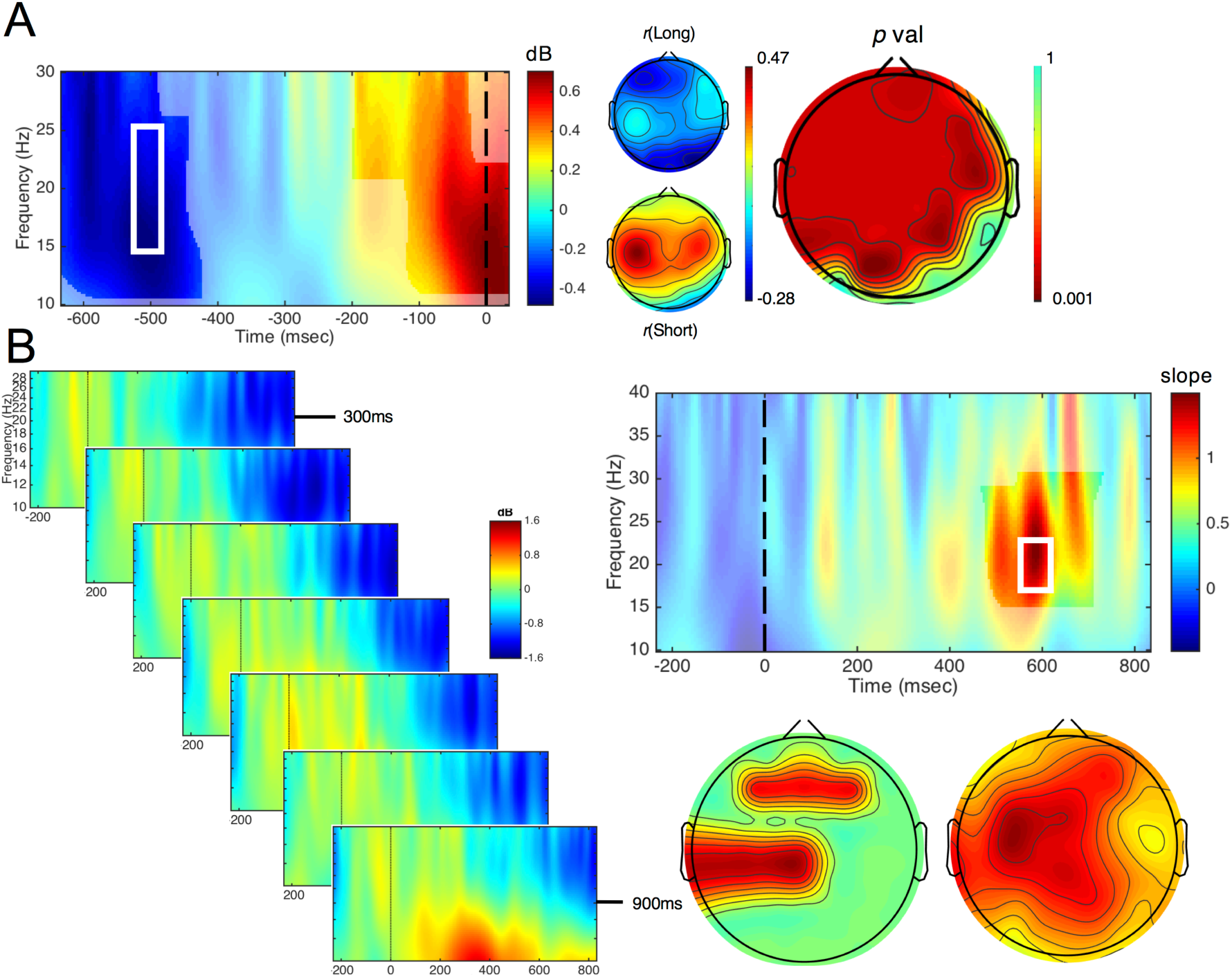
EEG Re-analysis of Wiener and Thompson (2015). **A**) Time/frequency plot of response-locked data, for the difference between long and short responses. Highlighted regions represent significant pixels from a cluster-correction procedure. A broad ERD/ERS profile was observed across alpha and beta frequencies, but was strongest within the beta range (electrode FCz). At right, the topographical distribution for long and short response choices and the significance across electrodes (cluster corrected), taken from within the white box at left. **B**) Time/frequency responses for onset-locked data, depending on the duration presented on the previous trial, collapsed across all durations presented on the current one (electrode FCz). Here, an increase in beta power was observed with longer durations. At right, slope values from a linear regression of power and duration revealed a significant cluster within beta frequency at 600ms. Below topographic plots display the distribution of slopes within the inset above (right) and their significance (left).

This finding suggests that beta oscillations were higher for trials on which the stimulus was classified into the long duration category. Yet, this pattern alone does not implicate beta oscillations in time perception, as the ERD/ERS pattern observed is commonly found when subjects prepare and then execute a motor response, regardless of the timing of that response (Meijer, te Woerd, & Praamstra, 2016; Kilavik, et al. 2013). As such, we next turned to the examination of so-called *direct* and *carryover* effects in these data (Aguirre, 2007). When spectral power was compared for each duration tested in our stimulus set, onset-related activity exhibited a stereotyped pattern of ERS at lower, alpha frequency between 200 and 400ms post-onset, followed by an ERD of higher, beta activity between 400 and 800ms post-onset. These windows correspond to the P300 and CNV components observed in the broadband data of the original study. Notably, for the direct effect of duration, no differences were detected at any time/frequency point between any of the tested durations, indicating that spectral power did not differ by the duration presented on each trial (supplementary figure 1). However, instead, we observed a significant cluster in the carryover analysis, by collapsing across all of the present trial durations and only observing how the data changed based on the prior trial duration. Here, we observed a positive cluster that peaked at ∼600ms between 17 and 23 Hz, within the beta frequency band in a frontocentral cluster. This cluster indicated that the spectral power increased as the prior trial duration increased (Figure 3b).

The finding that beta power was modulated by the prior, but not present trial duration, follows the CNV findings of the original study. In this work, the CNV also covaried linearly with the prior trial duration, with a higher amplitude for longer prior trial durations. In that study, this shift in the CNV was tied to fluctuations in the categorical boundary for classifying durations as long or short; a higher CNV amplitude for longer prior durations was associated with an increased probability of subsequently classifying the present duration as short. This finding was interpreted as the categorical boundary being shifted towards the value of the prior trial duration (Wiener, et al., 2014). Yet, the CNV signal here was also modulated by the present trial duration. The present findings with beta suggest that beta power is associated with the current value of the categorical boundary, or bisection point.

To further examine time/frequency effects, we next re-analyzed data from a simultaneous TMS-EEG study while subjects performed a visual temporal discrimination task (Wiener, et al. 2012). In this second re-analysis, we compared spectral changes resulting from stimulation of either the rSMG or midline occipital cortex (Mid-Occ) prior to the onset of a 600ms standard stimulus, which subjects were required to encode and subsequently judge against a comparison stimulus. In our original study, we found that stimulation of the rSMG, but not Mid-Occ, increased the CNV amplitude, and that the size of this effect correlated with an increased likelihood to judge the comparison stimulus as shorter than the standard; in other words, the perceived duration of the standard stimulus increased. When comparing rSMG and Mid-Occ stimulation conditions, we observed an increase in spectral power for a cluster spanning 300 to 500ms post stimulus onset between 25 and 27Hz, within the high beta frequency range (Figure 4). The window of this effect again corresponds to the CNV signal observed in the original study, and follows from the above re-analysis that an increase in beta frequency power follows an increase in judging the stimulus as long.

**Figure 4.**
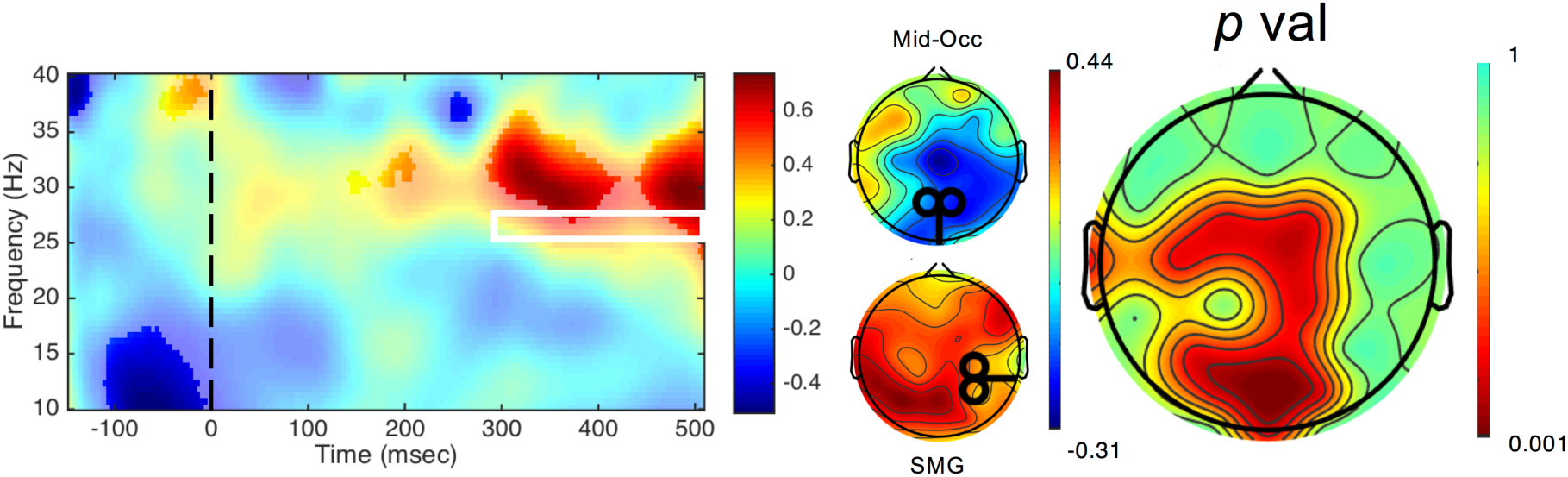
EEG Re-analysis of Wiener, et al. (2012). At left, time/frequency representation of the difference between rSMG and Mid-Occ stimulation revealed an increase in high beta power following rSMG stimulation (electrode Cz; highlighted regions are significant at p<0.05, uncorrected for visualization purposes). At middle, the distribution of beta power across electrodes for rSMG and Mid-Occ stimulation, from within the white box at inset, representing the cluster-corrected region of signifiance. At right, the significance of the difference within this region, across electrodes (cluster corrected).

Altogether, our findings suggest that the power of beta frequency oscillations contributes to the value of a remembered duration. In the re-analysis of Wiener and Thompson’s (2015) data, beta covariance with the prior trial duration suggests that this signal indexes the criterion duration being used for *comparison* to the present trial duration. The re-analysis of Wiener and colleagues’ (2012) data, in turn, suggests that beta power is modulated by rSMG stimulation while subjects are encoding a duration into memory that will subsequently be compared to another duration. In either case, we note that *higher* beta power is associated with *longer* remembered durations.

### tACS – Simulation

We analyzed simulation data of the electrical field induced by tACS at our electrode locations (FC1 & FC2). Using a finite element head model of a canonical brain, we observed a maximal normalized electrical field at the location of the SMA (Figure 1c). The effect covered both hemispheres, suggesting the effect of stimulation did not result from a lateralized impact. *tACS – Behavioral Analysis*

We next applied tACS at beta and alpha frequency bands in a new group of subjects (n=19) during a visual temporal bisection task. All subjects performed the task well, with no difference between stimulation conditions on the ability to perform the task (supplementary figure 2); a multivariate ANOVA of post-stimulation questionnaires did not reveal any significant differences between alpha and beta tACS on any of our measures, including tingling sensations and phosphenes (all *p* > 0.05; supplementary material). Here, psychometric and chronometric curves exhibited well-described effects (Kopec & Brody, 2010; Wiener, et al. 2014), with an increasing probability of classifying a stimulus as long with increasing duration, and a corresponding decrease in reaction times for longer stimuli (Figure 5). Notably, the bisection point (BP) for subjects in our sample was located above values typically found (Kopec & Brody, 2010), at a value above the arithmetic mean of the stimulus set [mean BP: 0.59 ± 0.027]; no difference between baseline BPs were observed [Wilcoxon Signed Rank Test: two-tailed *z* = −1.127, *p* = 0.278, based on 10,000 Monte Carlo simulations]. For chronometric data, a repeated measures ANOVA with frequency (alpha,beta), stimulation (baseline, concurrent), and duration (seven, log-spaced intervals) as within-subject factors revealed a significant main effect of stimulation [*F*(1,18)=21.447, *p*<0.001, η^2^_p_ = 0.54], but no interaction with the frequency of stimulation [*F*(1,18)=0.267, *p* = 0.612]. This effect was characterized by faster reaction times for both stimulation frequencies, with no difference in the size of the increase. For psychometric data, we conducted separate repeated measures ANCOVAs on BP and CV values, including frequency and stimulation as within-subjects factors, as well as mean trial count as a covariate (supplementary materials). For BP values, we detected no main effect of frequency [*F*(1,17)=0.992, *p* = 0.333] or stimulation [*F*(1,17)=0.535, *p* = 0.475], but did observe a significant interaction [*F*(1,17)=6.527, *p* = 0.021, η^2^_p_ = 0.277]. No significant effects were observed for a similar analysis of CV values (all *p* > 0.05). Post-hoc Signed Rank Tests revealed that beta, but not alpha, stimulation led to a significant decrease in the BP value [alpha: *z* = −1.408, *p* = 0.165; beta: z = −2.857, *p* = 0.003, Cohen’s *d* = −0.463], indicating a greater propensity to classify duration stimuli as “long” when receiving beta stimulation compared to baseline.

**Figure 5.**
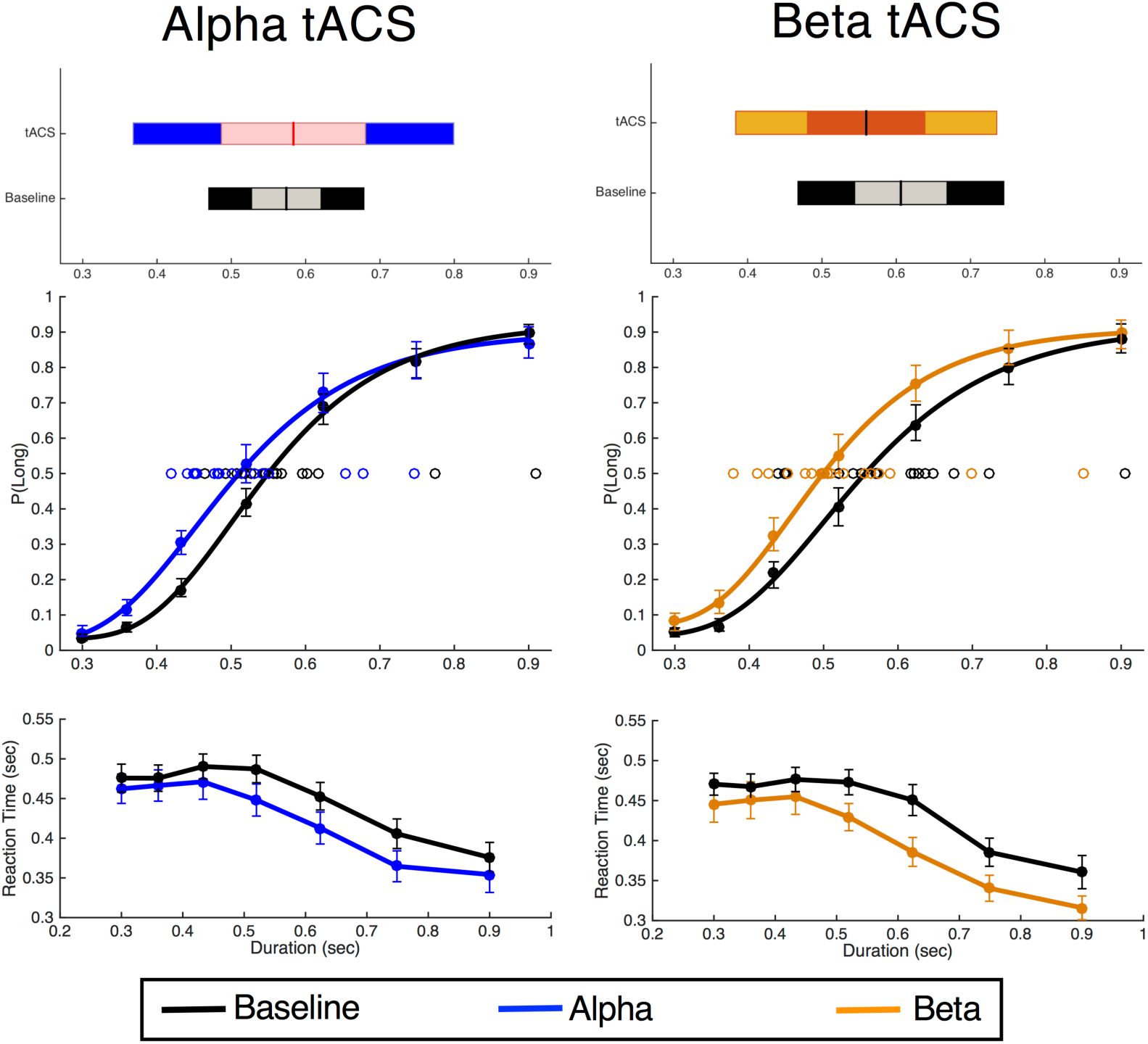
Results of tACS on temporal bisection. Top and middle graphs feature psychometric data, with the average proportion at which each interval was classified as “long”. Top graphs display the mean and spread of bisection points; inner bars and outer bars represent 95% confidence intervals and the standard deviation, respectively. bottom graphs feature chronometric functions, with the average reaction time for each interval, regardless of choice. Gumbel distribution curves for psychometric data are fit to the average proportions for visualization purposes only. Individual open data points on top panels represent the bisection point for all subjects. Stimulation was found to reduce reaction time for both frequencies, whereas beta stimulation alone significantly shifted the bisection point leftward, characterized by a greater propensity to classify stimuli as “long”. Error bars represent +/− standard error.

Altogether, the behavioral data indicated an increase in the BP value for beta stimulation, with no change from alpha stimulation. This effect was also independent of any changes in the CV value across stimulation conditions. Further, we observed no differences in any measures of carryover from the previous trial, either decisional or perceptual (Wiener, et al. 2014), indicating that the effect of stimulation on the BP was not the result of a sequential effect of the prior trial duration or decision (all *p* > 0.05; supplementary figure 3).

### Drift Diffusion Modeling

In order to further isolate the cause of the shift in BP for beta stimulation, we decomposed choice and reaction time data with a drift diffusion model (Wiecki, et al. 2013; Balci & Simen, 2014). Model fits demonstrated that the best-fitting or “winning” model for our design was one in which frequency, stimulation, and duration were all included (supplementary figure 4). This improvement in model fit was consistent for hierarchical, non-hierarchical, and regression methods. Analysis of the Gelman-Rubin statistic for multiple chains revealed a value of 1.0016 ± 0.0061(SD), indicating good chain stability. Visual inspection of model chains confirmed this finding, by demonstrating symmetrical traces and distributions, along with low autocorrelation levels. Finally, we performed a posterior predictive check of the winning model (Kruschke, 2013) by simulating 500 datasets using the mean posterior estimates for each node. Comparison of these datasets with the original data demonstrated that the model was able to reproduce the pattern observed in the actual dataset (supplementary figure 5).

We initially attempted in our model to replicate findings from Balci and Simen (2014), in their original formulation of the TopDDM model, as applied to temporal bisection. This model predicted four distinct patterns for *a, v, t,* and *z* values across duration (see Methods). Observation of the model fit values demonstrated the expected patterns for all four parameters. Specifically, we observed that threshold (*a*) values decreased for duration values closer to the middle of the stimulus set, and increased for extreme duration values. Drift (*v*) values were observed as negative for durations lower than the mean of the stimulus set, and positive for values above it, increasing linearly. Non-decision time (*t*) values decreased linearly for longer duration values. Finally, we predicted that the starting point (*z*) parameter values would increase linearly with duration. While a repeated measures ANOVA of starting point values did indicate a linear contrast with duration [*F*(1,18)=83.701, *p*<0.001, η^2^_p_ = 0.823], we note that starting point values monotonically increased only until the middle of the stimulus set (0.52s), before decreasing with longer durations. This pattern, while not predicted, does match with earlier electrophysiological findings from the temporal bisection task, which indicate that subjects accumulate information only until the categorical boundary (bisection point) is reached (Wiener & Thompson, 2015).

Further comparison between stimulation conditions revealed a number of distinct findings for alpha and beta stimulation. First, we observed that the threshold parameter decreased for both alpha and beta stimulation [*F*(1,18)=10.563, *p*=0.004, η^2^_p_ = 0.37]; yet, while this decrease was noticeably larger for alpha stimulation, no interaction between these effects was observed [*F*(1,18)=2.441, *p*=0.136]. For the drift parameter, a main effect of stimulation was again observed, with the drift rate overall shifting upwards (closer to the “long” duration boundary) [*F*(1,18)=47.174, *p*<0.001, η^2^_p_ = 0.724], but again with no interaction between stimulation frequencies [*F*(1,18)=0.556, *p*=0.465]. For non-decision time, no main effect of stimulation was observed [*F*(1,18)=0.193, *p*=0.66] nor an interaction with frequency [*F*(1,18)=0.012, *p*=0.914]. Finally, for the starting point parameter, no main effect of stimulation was observed [*F*(1,18)=0.929, *p*=0.348]; however, an interaction between stimulation and frequency was found [*F*(1,18)=12.612, *p*=0.002, η^2^_p_ = 0.412], with the starting point increasing closer to the long duration boundary for beta [Wilcoxon *z* = −2.857, *p* = 0.002, Cohen’s *d* = −0.46], but not alpha [Wilcoxon *z* = −1.368, *p* = 0.182] stimulation.

**Figure 5.**
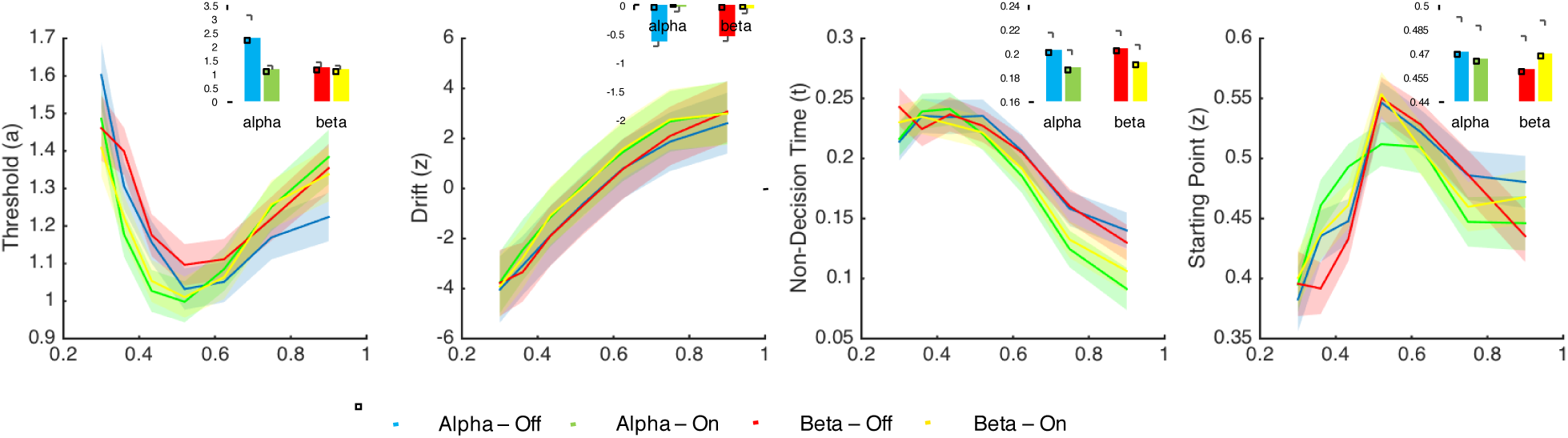
Fits of model parameters for the hierarchical drift diffusion model. Insets represent model parameters collapsed across duration for each stimulation condition. Shaded regions and error bars represent standard deviation across subjects. Model parameters confirm earlier findings (Balci & Simen, 2014) and show that stimulation decreases the threshold and increases the drift rate at both frequency ranges, but exclusively increases the starting point for beta stimulation. No impact on non-decision time was found.

While the results above highlighted a number of distinct effects for tACS, including a differential effect on the starting point parameter between alpha and beta stimulation, we note that the effects observed may have been driven by the method used for fitting our model to the data. Specifically, we initially employed a hierarchical design, in which the individual subject posterior estimates are constrained by the group distribution (Kruschke & Vanpaemel, 2015). As such, our findings may have been driven by the non-independent nature by which they were sampled. To assess this, we conducted a separate, non-hierarchical sampling procedure, wherein each subject was modeled separately, and individual parameters estimates were combined afterwards. Comparison of the non-hierarchical model to the hierarchical one demonstrated relatively good concordance between both fitting methods (supplementary figure 6a), with the same pattern observed between stimulation and frequency-of-stimulation conditions. The only exception was for the threshold parameter, in which the non-hierarchical method did not reveal a decrease in the threshold for alpha tACS, suggesting caution in interpreting this result, a point we turn to below.

Given the discrepancy between hierarchical and non-hierarchical methods with regards to the threshold parameter for alpha stimulation, we sought to apply a hierarchical regression method to our data. Specifically, the hierarchical method we employed utilized separate nodes for each condition and each subject, and does not take into account within-subject covariance across conditions. To provide greater specificity of our model, we utilized the HDDMRegressor class of the HDDM toolbox (Wiecki, et al. 2013) to construct a hierarchical linear model. To accomplish this, each model DDM parameter was described by a general linear model with a random intercept for each subject, frequency (alpha, beta) and stimulation (pre-stim, stim) conditions as dummy variables, and duration as a continuous covariate that interacted with frequency and stimulation.

The results of the regression analysis are displayed in supplementary figure 6b. In general, the regression findings accord with the hierarchical and non-hierarchical findings. For the threshold parameter, we observed that slope value for stimulation was negative and did not fall within a region of practical equivalence (ROPE; Kruschke, 2011) of [−0.1 – 0.1] (β= −0.4476, 95% Credible Interval [−0.5166, −0.3822]), indicating that threshold values decreased with stimulation, whereas the interaction effect between stimulation and frequency encompassed the ROPE (β= 0.0669 95% CrI [−0.0215, 0.1594]), indicating that threshold values decreased with stimulation, but this effect did not vary by stimulation frequency. For the drift parameter, the stimulation slope was positive and was not within the ROPE (β= 8.2038, 95% CrI [8.0231, 8.3818]), indicating that drift rates overall increased with stimulation; notably, the interaction effect between stimulation and frequency was also positive and was not within the ROPE (β= 0.3821, 95% CrI [0.1418, 0.6399]), indicating that the increase in drift values was differential between alpha and beta frequency. For non-decision time, the stimulation slope was negative and did fall within the ROPE (β= −0.0489, 95% CrI [−0.0537, −0.0384]), as did the interaction (β= −0.0156, 95% CrI [−0.0268, −0.01]) indicating that non-decision time decreased with stimulation, but this effect could not be reliably differentiated from a null effect. Finally, for the starting point parameter, the stimulation slope was negative and fell within the ROPE (β= −0.1373, 95% CrI [−0.2336, −0.04]), whereas the interaction effect was positive and did not fall within the ROPE (β= 0.2552, 95% CrI [0.1102, 0.4145]), indicating there was no main effect of stimulation, but that stimulation and frequency interacted.

Overall, the results of the regression analysis corroborated the findings of the hierarchical and non-hierarchical fitting methods. We choose to rely on findings that agree across all three fitting methods when describing our results. Specifically, tACS, regardless of frequency, led to a decrease in the threshold and an increase in the drift parameters. For the non-decision time parameter, no effect of stimulation or interaction with frequency was observed. Finally, for the starting point parameter, no main effect of stimulation was observed, but an interaction with frequency was found, wherein the starting point shifted positively, towards the long duration boundary, for beta stimulation.

## Discussion

Beta oscillations have emerged in the past few years as a candidate mechanism for coordinating timing functions in the human brain. These findings suggest that beta oscillations have a supramodal property, in that they cover timing and predictive abilities regardless of the task context or modality of timed stimuli. In order to determine what role, if any, beta plays, we conducted two re-analyses of previously-collected EEG data, followed by a novel tACS study that employed stimulation at two distinct frequency bands (beta and alpha) while subjects performed a temporal bisection task. The result of these analyses point to a specific role for beta oscillations in timing, which we now elaborate on further.

In re-examining previously collected EEG data from an auditory temporal bisection task (Wiener & Thompson, 2015), we first observed a curious finding: although oscillatory power, spanning a wide range that encompassed the beta band was modulated by the choice on each trial, with relatively lower beta power (desynchronization) prior to classifying a stimulus as “long” in duration, this effect was not parametrically modulated by the duration presented on each trial. Instead, beta power covaried linearly with the duration presented on the previous trial, with higher beta power for longer durations. One possible explanation for this finding is that beta power indexed the duration to which each presented value was being compared. That is, beta power serves as an index of the memory standard by which durations are categorized. Consider, the memory standard, as measured by the bisection point of the observed psychometric function, fluctuates on a trial-by-trial basis, and gravitates towards whatever duration was presented on the prior trial (Wiener, et al. 2014; Bausenhart, Bratzke, & Ulrich, 2016). As such, beta power may reflect here a shift in the value of the memory standard over time. From a Bayesian perspective, this may be interpreted as an iterative updating of the prior probability distribution. Consistent with this view, recent evidence suggests that beta oscillations may track the trial-by-trial weight of evidence for making decisions (Gould, et al. 2012).

In our second re-analysis, we examined the effects of rTMS on the rSMG of the parietal lobe while subjects performed a visual temporal discrimination task. In this task, the temporal referent that subjects must use for comparison is presented on every trial. Stimulation applied immediately prior to the onset of this standard stimulus was found to shift comparison responses in a manner to suggest that subjects encoded the standard stimulus as longer in duration than it actually was (Wiener, et al. 2012). The original analysis also found that the frontocentral CNV signal during the encoding of the standard stimulus increased with stimulation. Similarly, we observed an increase in high beta power during the encoding of stimulus duration. The direction of this effect is consistent with what we observed in our previous re-analysis – increased beta power was associated with a longer remembered duration. We reconcile this finding with the re-analysis of the previous study by noting that, here, subjects were encoding the standard stimulus duration into memory. As such, the observed increase in beta power may reflect a longer encoded duration. Similar findings have been observed in the interval timing literature in animals, where pharmacological manipulation alters remembered duration via a hypothetical memory translation constant (Meck, 1996), invoked as a part of scalar timing theory (Gibbon, et al. 1984). Altogether, both re-analyses established a supramodal role for beta oscillations in time perception, that may be linked to the retention of a memory standard for duration, rather than the perceived duration itself.

While the above re-analyses are suggestive of the involvement of beta oscillations in perceptual timing, they do not reveal a causal link between beta power and timing functions. To establish this, we applied tACS at two distinct frequencies during a visual temporal bisection task. Here, we first observed that stimulation at both alpha and beta frequencies led to a decrease in reaction time functions, such that subjects were faster at making responses when receiving stimulation compared to when not. Although it may be tempting to state that stimulation made subjects faster, it should be noted that the pre-post design of our study leaves open the possibility that subjects were faster due to practice with the task; stimulation always occurred after the baseline task (Figure 1), and so subjects may have improved simply by familiarity with the task demands. A second observation was that the bisection point (BP) significantly shifted leftward for beta, but not alpha stimulation. In this case, subjects became more likely to classify stimuli as “long” in duration. Notably, this effect was independent of any carryover effect from the previous trial or decisional bias from the previous choice (Wiener, et al. 2014).

In order to further disentangle the possible bases for this effect, we decomposed choice and reaction time data with a drift diffusion model inspired by recent work (TopDDM; Balci & Simen, 2014). Across several methods of fitting the data to individual subjects, three findings were apparent. First, the threshold boundary, which the decision variable must cross in order for a response choice to be made, was lower for both alpha and beta stimulation regimes, indicating that subjects were less cautious, requiring less evidence, before making a choice (Voss, Rothermund, & Voss, 2004). Second, the drift parameter was significantly higher overall for both alpha and beta stimulation. In this case, a “higher” drift parameter indicates that the decision variable drifted more quickly towards the long duration boundary than the short duration one. This is due to the way in which we parameterized drift: negative drift values indicate an accumulation towards the short duration boundary, whereas positive values indicate accumulation to the long duration boundary (cf. Balci & Simen, 2014). Finally, the starting point parameter moved closer to the long duration boundary, but only for beta, and not alpha stimulation. For many formulations of the drift diffusion model, a change in the starting point indicates a shift in bias, such that subjects favor one response over another and so require less evidence before committing to a decision in that direction. Here, a shift in the starting point carries a different interpretation; specifically, the starting point is meant to reflect the value of the first-stage timing process *A* at stimulus offset (Figure 2). A higher starting point thus reflects a larger subjective value of time. We note that it is unlikely that the effects observed were due to a simple shift in bias; if that had occurred, we would likely also have seen a subsequent impact on the carryover effect of the previous decision (Wiener, et al. 2014), which was not observed.

### Functional consequence of beta and alpha manipulation

The results of the drift diffusion modeling suggest that beta oscillations exclusively shift the starting point of the decision process in temporal bisection, an effect that may be tied to a change in the first-stage temporal accumulator value. Yet, we note that a change in this effect does not match well with the observed findings. At first glance, an increase in the temporal readout value used for measuring subjective time would be expected to make subjects classify stimuli as long more often. However, the temporal bisection task requires subjects to continuously compare each duration to a running average of the durations they previously experienced. As such, every longer-perceived duration would in turn go into the memory used for judgment on the next trial. As such, any effect would be short-lived, as subjects shifted to a new baseline, consistent with so-called “clock pattern” effects observed for pharmacological agents affecting the dopamine system (Meck, 1996). Further, the results of our EEG re-analyses did not suggest a role of beta oscillations in subjective time readout, as evidenced by a lack of effect for beta power while subjects measured stimuli of different lengths. Instead, these findings suggest a role for beta in the memory process, specifically for the encoding of durations into memory. Here, the effect of stimulation would be to shift the boundary used for comparing durations, rather than the perception of any individual duration itself. Here, the effect would be comparable to a “memory pattern” effect, in which the psychophysical function shifts away from baseline and remains separate, as accurately perceived intervals are subsequently entered into memory as a different value. To disentangle which pattern exists in the data, we conducted a subsequent sliding-window analysis of the data. Here, we used a sliding window of 131 trials – the minimum needed for a sufficient trial count for each duration to accurately fit the data – and continuously analyzed the bisection point over the course of the stimulation session. The results are presented in supplementary figure 7, and show that, contrary to a clock pattern, the effect was maintained throughout the session, consistent with a memory pattern.

To reconcile the memory pattern we observed with the drift diffusion model findings, we suggest that the starting point, rather than representing a subjective time estimate at offset, instead reflects in-part the location of the categorical boundary. Under this framework, if the categorical boundary is shifted lower, then the distribution of intervals in the stimulus set becomes uneven, with more long intervals than short ones. As such, the starting point will shift towards the long duration boundary because long durations are more likely to occur. Further testing will be necessary to see if this bears out in behavioral data, by testing subjects with different distributions of intervals that are unevenly skewed towards the longest or shortest durations in the stimulus set, a difference that has been shown to reliably shift the bisection point (Brown, et al. 2005).

The lack of a consistent effect for alpha stimulation also presents an interesting finding. Although we did not find any effect in the alpha range in our EEG re-analyses, alpha oscillations have previously been associated with time estimation, particularly for visual stimuli (Wiener & Kanai, 2016). Notably, although no significant effects were observed in mean differences across subjects, a closer examination of the distribution of performance with alpha stimulation reveals an interesting finding. Specifically, during alpha stimulation, we observed that the bisection point values became more heterogeneous between subjects than prior to stimulation. This observation was confirmed with a simple two-sample *F* test^1^ comparing the variances of stimulation and pre-stimulation conditions, revealing that the distribution of bisection points became more variable across subjects during stimulation [*F*(18,18) = 4.273, *p* = 0.0035]; no difference in variances was observed for beta stimulation [*F*(18,18) = 1.608, *p* = 0.3221]. This finding suggests that alpha stimulation may have had some impact, but one that was inconsistent between subjects. One possible explanation for this effect is the finding that alpha stimulation can demonstrably impact individual alpha frequency (IAF; Cecere, et al., 2015). More specifically, IAF is thought to reflect the intrinsic resting “peak” of alpha oscillations in an individual subject. Recent work has found that IAF also correlates with perceived visual timing (Samaha & Postle, 2015; Babiloni, et al. 2004), and that manipulation of this rate can reliably shift the perceived timing of visual events (Cecere, et al. 2015). However, much of this work relies on using a baseline measure of IAF in each subject, and then stimulating each individual at that rate (Zaehle, Rach, & Hermann, 2010). In our case, all subjects were stimulated at the same rate (10Hz), and so some subjects may have had their IAF speed up and other slow down. If the speed of intrinsic alpha oscillations determines in part the timing of visual stimuli, it’s possible that alpha stimulation would induce clock pattern type effects, but without titrating stimulation frequency to IAF, this effect would wash out, potentially leading to the increase in between-subject variability we observed. By using a baseline measure of IAF, this effect could be explored with further tACS.

### Anatomical locus of beta stimulation

In the present study, tACS was administered frontocentrally, over electrodes FC1 and FC2. Electric field modeling in a simulated subject suggests the impact of this montage is maximal within the confines of the supplementary motor area (SMA). We chose this site on the basis of functional evidence suggesting the SMA serves as part of a core network mediating time perception (Wiener, et al. 2010). Indeed, numerous studies now demonstrate SMA activity while subjects are engaged in a timing task (Schwartze, Rothermich, & Kotz, 2012). With regard to the precise function of this region, current debate exists regarding its role in timing (Coull, Vidal & Burle, 2016; Kononowicz & Penney, 2016). One particular theory is that the SMA reflects a temporal accumulator process, used for measuring elapsed durations (Casini & Vidal, 2011). In this case, the SMA would correspond to the first-stage drift process of TopDDM. Yet, as observed, the shift in the bisection point function for beta stimulation is unlikely to be driven by a clock pattern effect. A second theory is that SMA activity reflects the memory criterion for making temporal decisions (Ng, et al. 2011; Wiener & Thompson, 2015). Given the pattern of results described above, we suggest this to be the more likely function of the SMA for timing.

Outside of the timing literature, beta oscillations have previously been associated with the SMA with a variety of functions. Indeed, beta-band oscillations have traditionally been associated with motor functions (Baker, 2007), and although the present study required a speeded response, beta activity covaried with the duration presented on the previous trial, not the present one, suggesting an independence from motor processes. Neural oscillations have been suggested as an organizational framework for neuronal networks, whereby activity may be synchronized across neural ensembles (Lakatos, et al. 2008). In this regard, beta oscillations may be less well understood than oscillations in other frequency bands (Engel & Fries, 2010). However, recent research has implicated frontocentral beta oscillations in numerous non-motor functions, such as perceptual decision-making (Haegens, et al. 2011) and local visual feature processing (Romei, et al. 2011). Yet, this does not preclude the involvement of the motor system in non-motor functions. Indeed, the SMA and basal ganglia, likely generators of the CNV and beta oscillations observed in the present study (Praamstra, et al. 2006; Fan, et al. 2007; Bartolo, et al. 2014), have been implicated in temporal perception, regardless of movement requirements (Wiener, et al. 2010). Importantly, we here distinguish the beta oscillations observed from other beta oscillations in the brain. That is, given the anatomical locus of our effect was confined to the SMA, we likely impacted beta oscillations within the cortico-striato-thalamic network. In contrast, beta oscillations have also been observed at occipital and parietal sites, with beta power in this region suggested to underlie visual excitability (Samaha, Gosseries, & Postle, 2017). We suggest it unlikely that our stimulation paradigm impacted visual experience directly, as subjects did not report a higher number of phosphenes or other visual phenomenon when receiving beta stimulation over alpha stimulation.

Another noteworthy aspect of our study is that stimulation effects of the SMA on timing are rare (Wiener, 2014). Indeed, the SMA has been stimulated in numerous paradigms with transcranial magnetic stimulation, but only two studies that we are aware of have demonstrated an effect (Dusek, et al. 2011; Mendez, et al. 2017). Both studies employed continuous theta burst stimulation, a method that has been suggested to induce a stronger and longer-lasting effect than conventional, repetitive TMS paradigms. However, in both cases, the effects observed were either weak (Dusek, et al. 2011) or indistinguishable from stimulation at other sites (Mendez, et al. 2017). One study that we are aware of has stimulated the SMA with tES (Dormal, et al. 2016). Here, the authors found that transcranial random noise stimulation (tRNS), a method in which the stimulation frequency varies randomly between 0.1 and 640 Hz, had no impact on timing when administered over the SMA. Given that our findings point to a specific role of beta frequency stimulation, one possible explanation for these discrepancies is that SMA effects are frequency dependent. As such, stimulation of this region is more likely to lead to an effect if the paradigm includes a dominant frequency within the beta band.

### Physiological locus of beta stimulation

The results of our re-analysis of two EEG datasets, as well as our stimulation findings, provide an overall role for beta oscillations in temporal memory. Moreover, these findings suggest a linear relationship between memory and beta power, with relative increases in beta power associated with longer retained duration memories. This finding is consistent with previous work, demonstrating a relationship between beta power and timing (Wiener & Kanai, 2016). The closest analogue to our findings is a re-analysis by Kononowicz and van Rijn (2015), in which higher beta power was associated with a longer produced duration. In this study, subjects were required to press a button when a prescribed duration had elapsed (2.5-seconds). Here, according to scalar timing theory, a produced interval will be randomly longer if either the temporal accumulation rate is high, or if the retrieved memory used for a criterion is long. Our findings suggest it is the latter case that explains the association between high beta power and longer productions.

Yet, our findings also present another curiosity – if higher beta power is associated with longer duration memories, then in order to explain our results, tACS must have *reduced* beta power. tACS is an effective means for entraining oscillations at a particular frequency band, but whether the effects raised or lowered power in the current study cannot be demonstrated. Yet, we note that the impact of tACS on oscillatory power is highly state-dependent (Feurra, et al. 2013; Schmidt, et al. 2014; Helfrich, et al. 2014). Feurra and colleagues (2013) demonstrated that the effect of beta tACS was dependent on the motor imagery a subject was engaged with at the time of stimulation. In our study, stimulation effects likely interacted with the active necessity of the subject to maintain vigilance. Additional work with simultaneous tACS-EEG will be necessary to determine the impact of stimulation on beta power.

With regards to the physiological basis of our effect, we note that beta oscillations, particular in the motor circuit, are highly associated with dopamine and cholinergic function (Kondabolu, et al. 2016; Jenkinson & Brown, 2011). Both neurotransmitter systems have been implicated in timing functions (Coull, Cheng, & Meck, 2011). Indeed, dopamine and acetylcholine manipulation in animals are capable of inducing clock and memory patterns in timing behavior, respectively. One particular impact of beta stimulation may have been to alter the dynamic balance between these two neurotransmitter systems, thus inducing the change in performance we observed. Whether or not beta oscillations observed with EEG or impacted with tACS change, depending on the dopamine state (Coull, et al. 2011) or effective level of acetylcholine (Meck, 1987), remains to be seen.

### Conclusions

The results of our two re-analyses and subsequent tACS study together provide a causal linkage between beta oscillations and time perception. These findings suggest that beta power contributes to the duration of an interval in memory, which impacts how a subject will judge future intervals. Further, our results are unlikely to be due to alternative effects on motor preparation or decision bias, and instead reflect an intrinsic function of beta oscillations. These findings may be connected to future work on neural oscillations and brain stimulation in time estimation. Further, it leaves open the possibility of other frequency bands not explored here, such as delta and theta frequency, or high gamma frequency, each of which may be involved in their own functions for timing.

## Supplementary Material

**Supplementary Figure 1.**
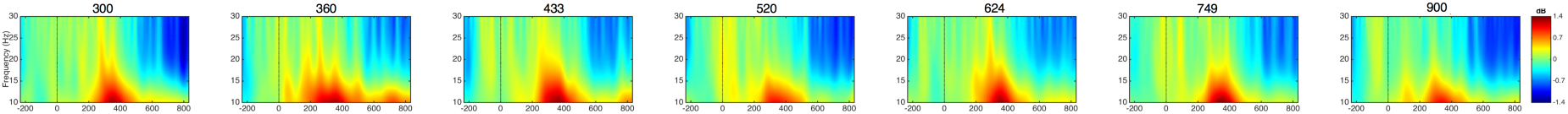
Direct effects re-analysis from Wiener & Thompson (2015). Time/frequency plots of onset-locked responses to each of the seven durations presented on the current trial (electrode FCz). No significant differences at any time/frequency point were observed.

**Supplementary Figure 2.**
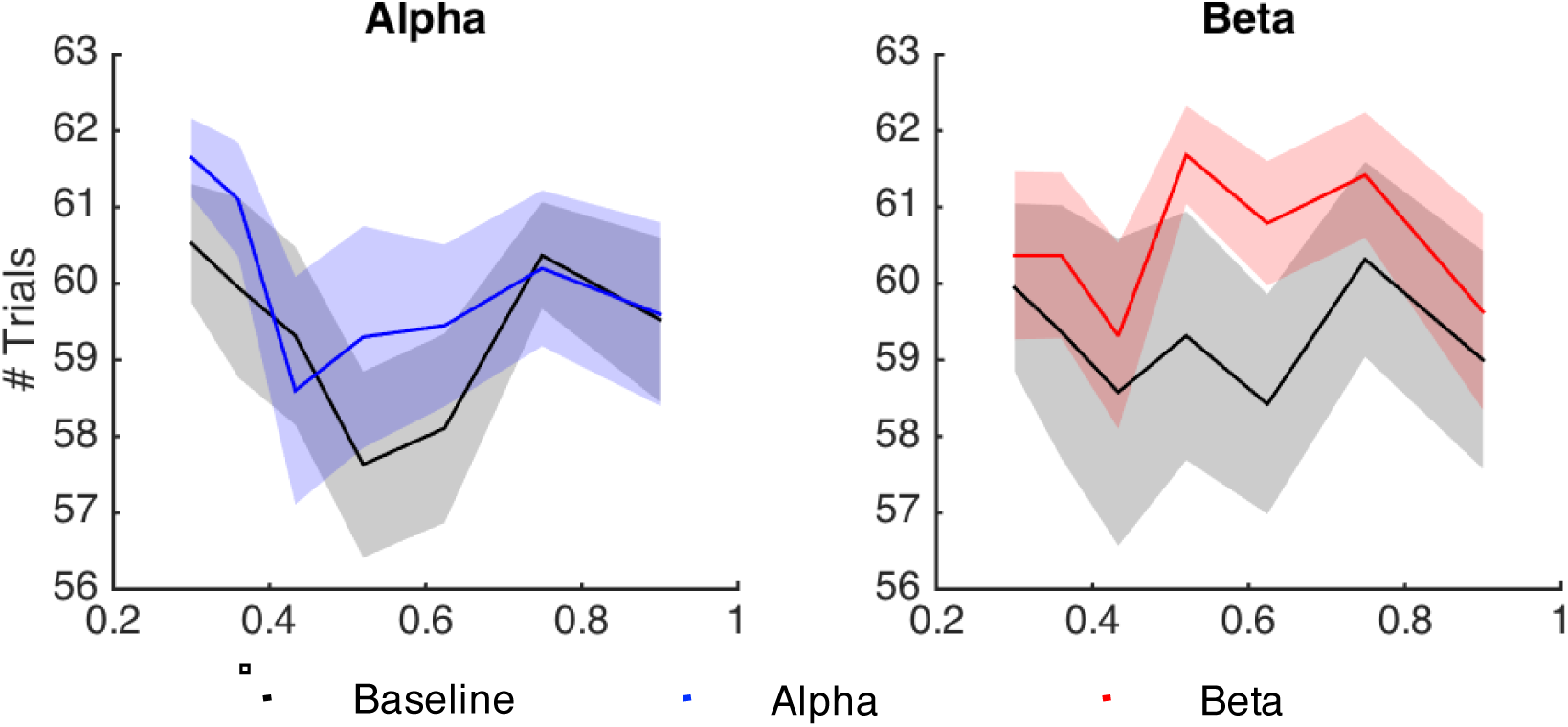
Number of trials for each interval across stimulation conditions after removal of trials where the reaction time exceeded 1000ms. A repeated measures ANOVA found a significant main effect of stimulation [*F*(1,18) = 5.107, p = 0.036, η^2^_p_ = 0.221], and a frequency by duration interaction [*F*(1,18) = 2.917, p = 0.021, η^2^_p_ = 0.139]. Specifically, fewer trials were filtered out for the beta stimulation session than for the alpha stimulation session. Due to these differences in trial count, we included the average number of trials per subject as a covariate in subsequent analyses of the bisection point, where the number of trials can impact the shape of the distribution (Fründ, et al. 2011). Shaded regions represent +/− standard error.

**Supplementary Figure 3.**
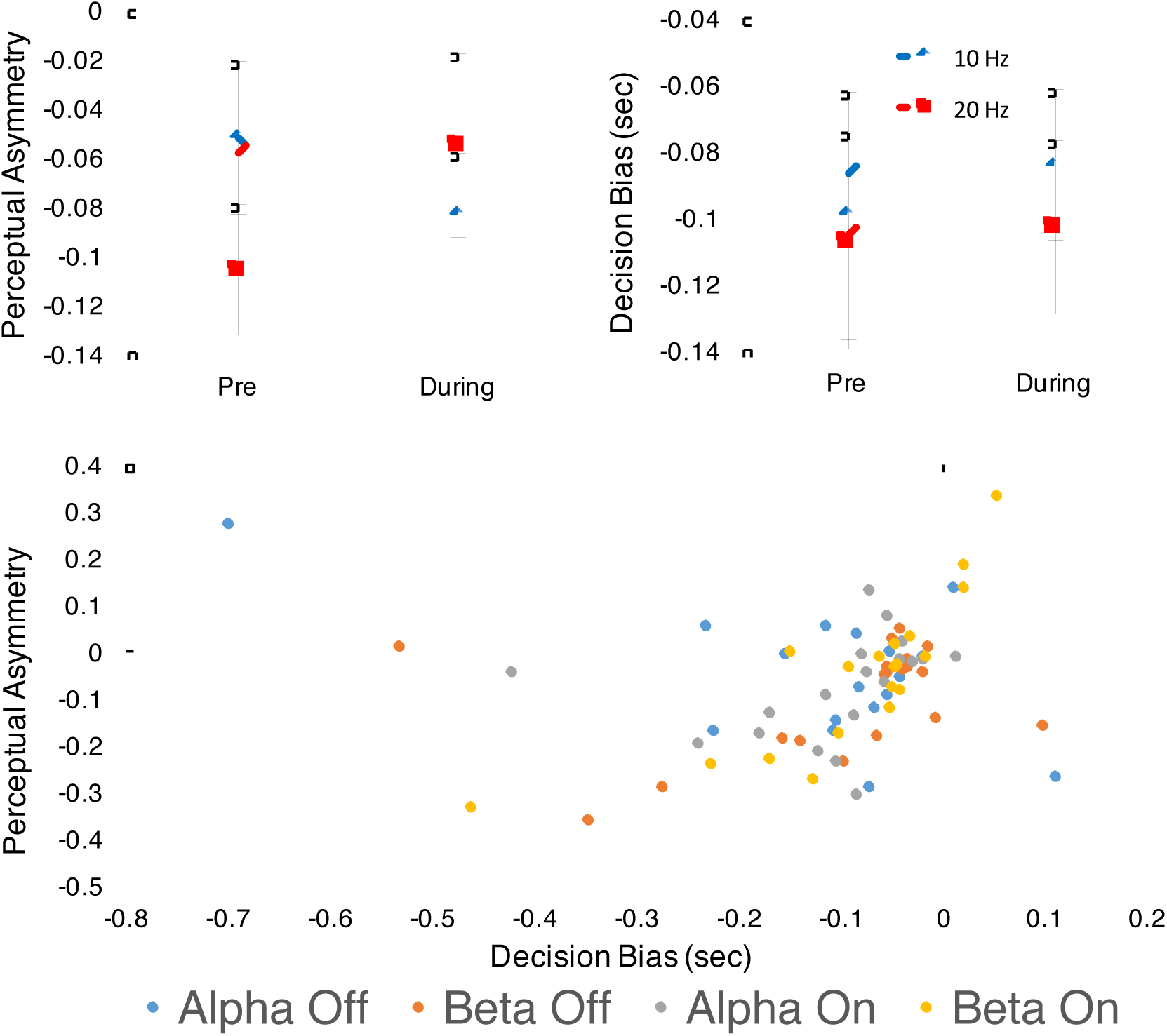
Carryover analysis of psychometric data (Wiener, et al. 2014). Top graphs display the mean perceptual asymmetry and decision bias values, respectively. Perceptual asymmetry reflects the influence of the preceding trial duration on the present trial duration; negative values indicate assimilation whereas positive values indicate repulsion. Decision bias reflects the influence of the preceding trial decision on the present trial decision; negative values indicate subjects are more likely to respond on each trial with the response made on the previous one. Bottom scatterplot displayed individual subject data points, demonstrating a correlation between both types of carryover, replicating prior findings (Wiener, et al. 2014). No significant effect of stimulation or frequency was observed for any carryover measure. We note that the removal of outliers from the above graph did not change the overall findings.

**Supplementary Figure 4.**
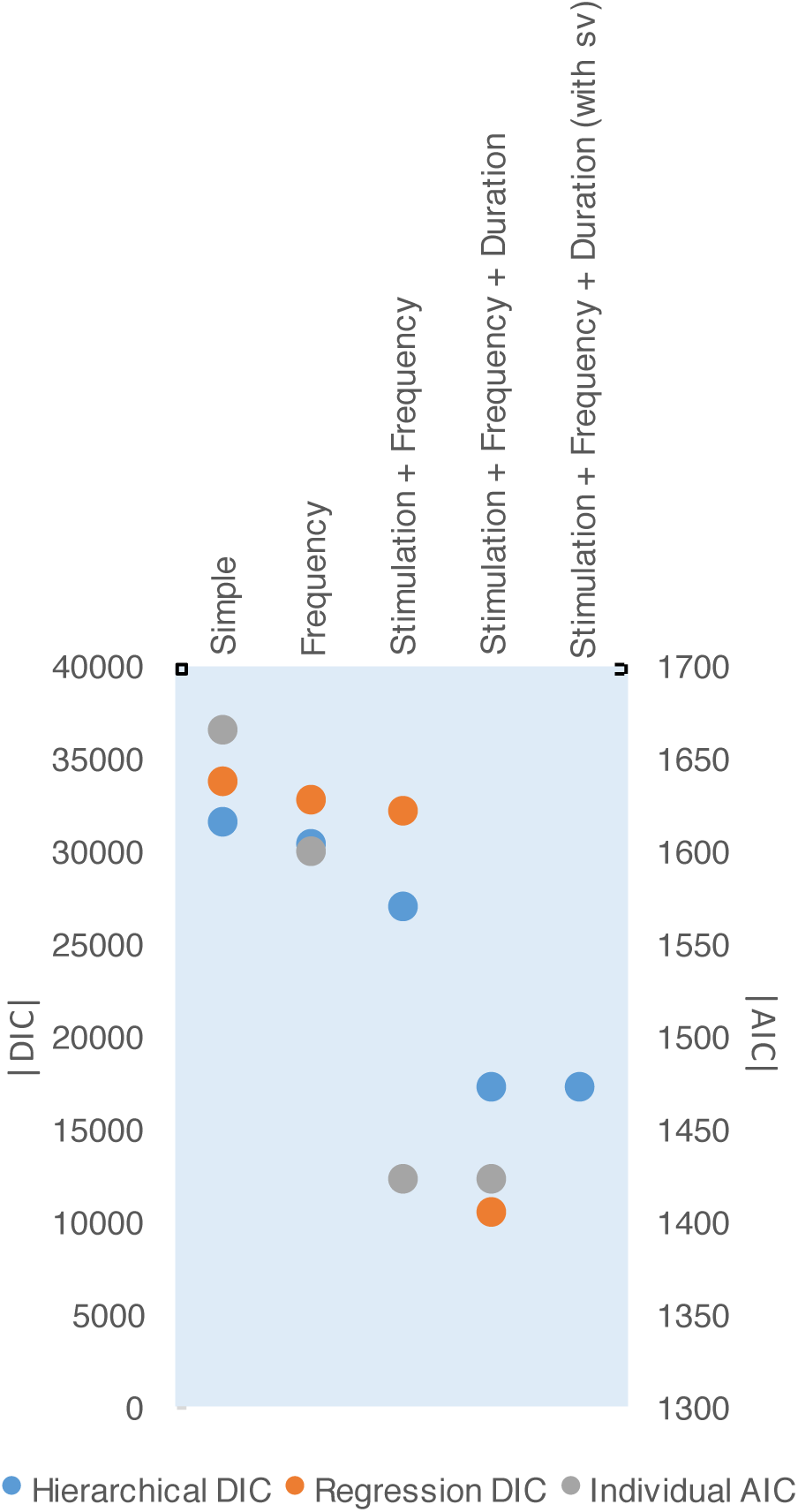
Fitting results of the different models used. DIC values are used for hierarchical and regression models, whereas the average AIC value is used for non-hierarchical fitting. In all cases, improved fits were found with more complex models, with the best fit for the Stimulation + Frequency + Duration model. A fifth model was also tested hierarchically, in which the variability of the drift rate (sv) was allowed to vary across duration (Balci & Simen, 2014). However, this model did not improve the fit over the best fitting model (ΔDIC <10).

**Supplementary Figure 5.**
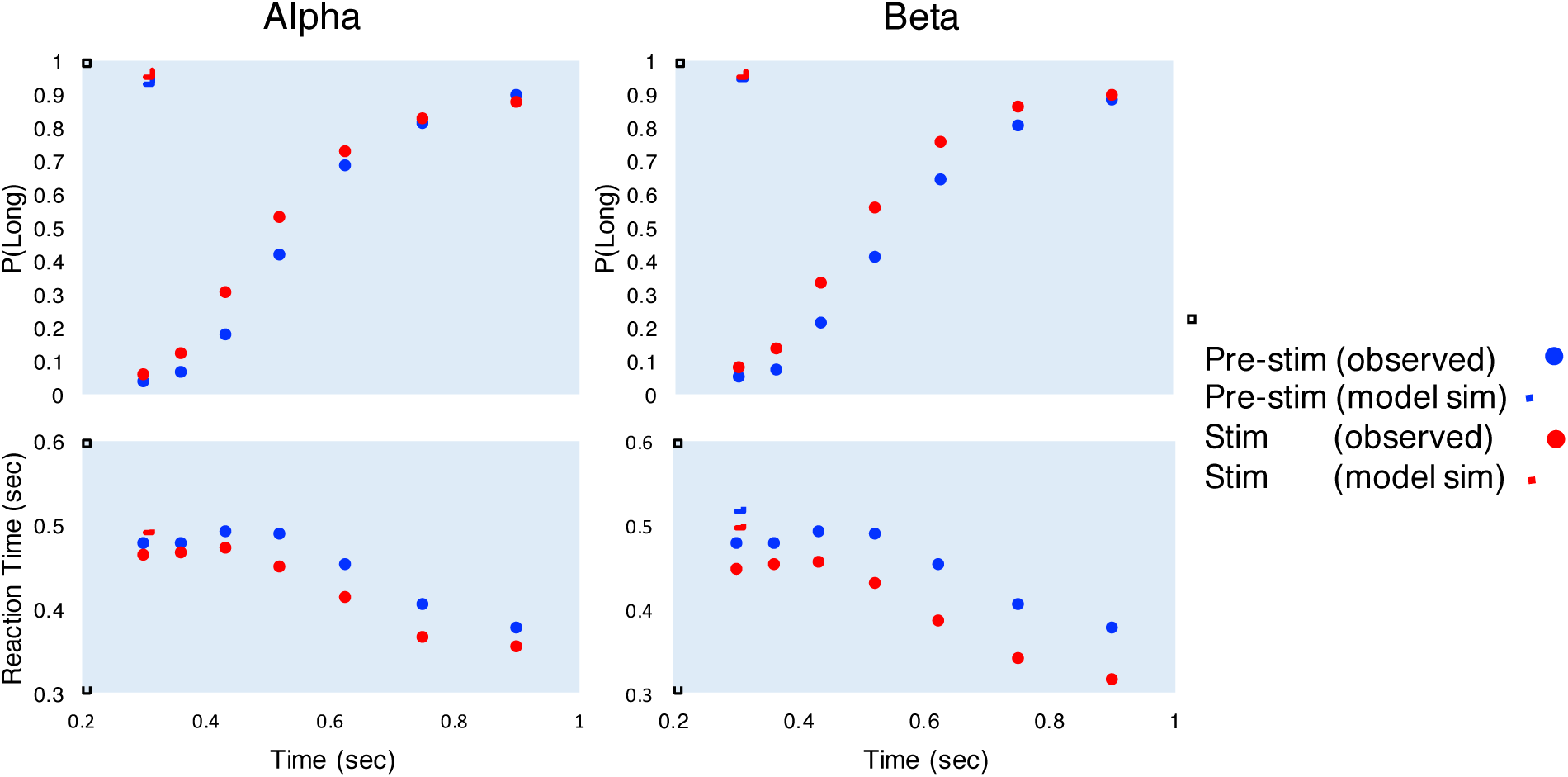
Results of posterior predictive checks. Model simulations were run, using the peak parameters from the hierarchical DDM, to generate 500 “subjects”. Average model simulated values are plotted here for psychometric and chronometric data (filled lines) over average subject performance (closed points). In general, model simulations produced the same observed pattern in the observed data, with faster RTs for stimulation conditions, and a more pronounced leftward shift for beta stimulation.

**Supplementary Figure 6.**
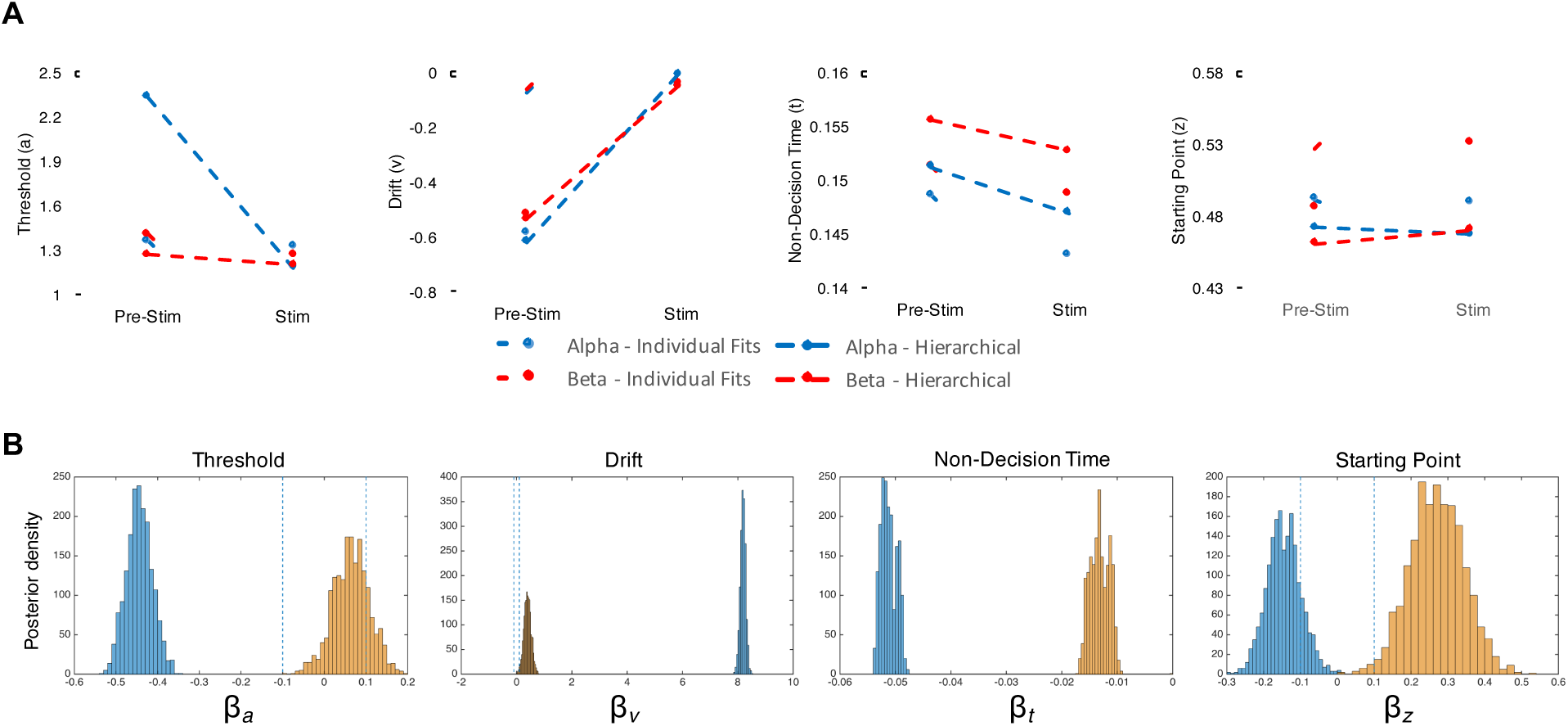
Model fitting results summary. A) Results for hierarchical and non-hierarchical fitting methods, collapsed across duration, for all four parameters in the model (*a,v,t,z*). In general, individual and hierarchical fits agreed. The only exception is for threshold values, where individual fits suggest an interaction between stimulation and threshold. For the starting point, although pre-stimulation values differ, individual and hierarchical methods both suggest an interaction. B) Results of the regression analysis. Displayed are the posterior probability density distributions for each parameter value. Blue distributions represent the main effect of stimulation, whereas gold distributions present the interaction between stimulation and frequency. Dashed lines indicate the ROPE; distributions where the highest density region (95% credible interval) falls within this region cannot be reliably distinguished from a null effect. For non-decision time, both distributions fall within the ROPE.

**Supplementary Figure 7.**
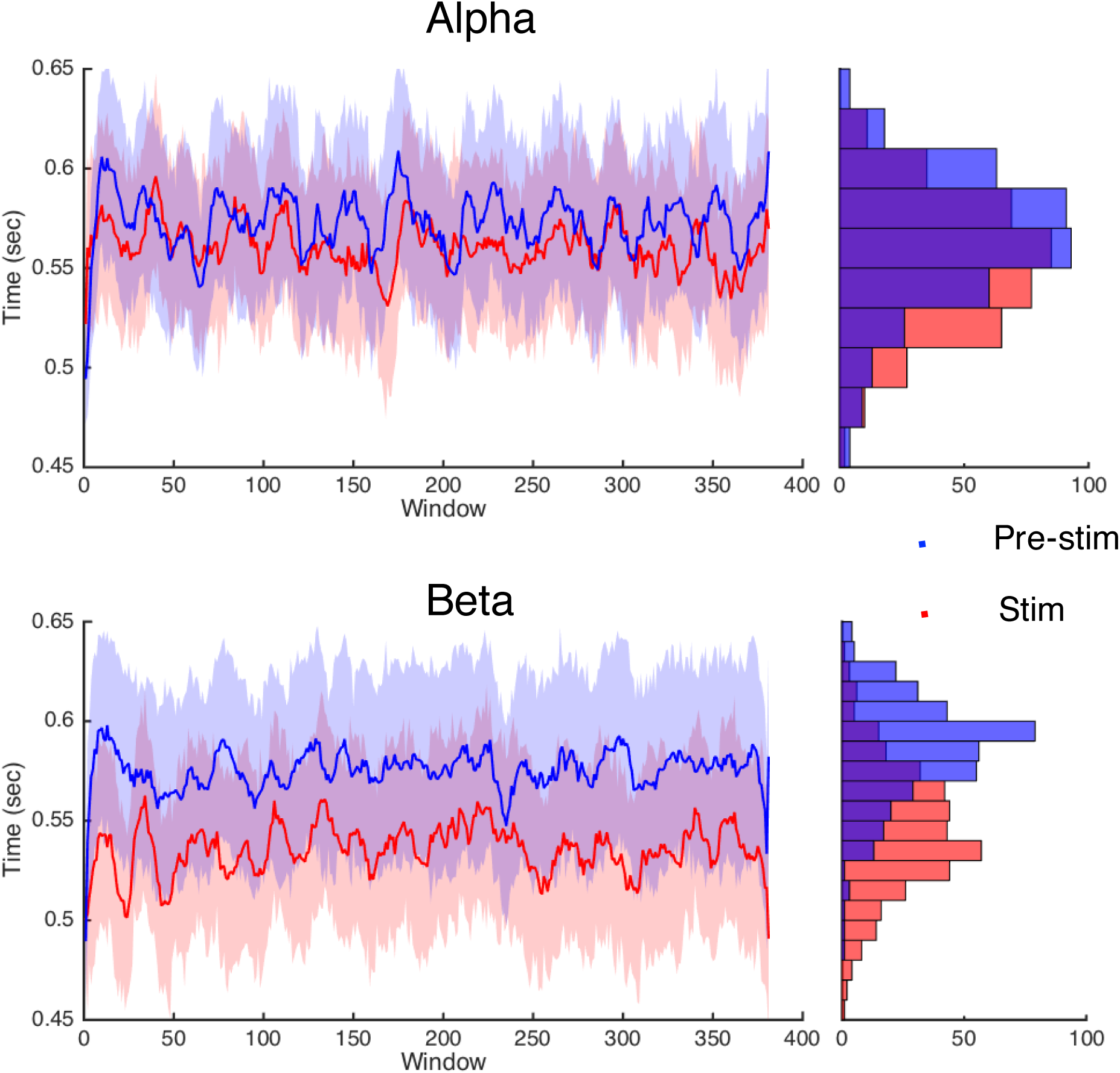
Sliding window analysis of within-session effects. A sliding window of 131 trials was ran across choice data for each session and the bisection point was calculated within each window as described in the methods. This window was chosen as the minimum size needed for each of the seven durations to have the minimum number of trials needed to fit a psychometric function (n=8). Data displayed above are smoothed by a 10-point moving average for the visualization of within-session trends. At right, frequency distributions of the bisection point values for each trace. The overall finding of above is that the shift in the bisection point values for beta stimulation are largely consistent across the session, rather than larger at one point in time over another. Shaded regions represent +/− standard error.

**Supplementary Table 1.** Sensation questionnaire responses during and after tACS. Bold points represent the average score, with italicized scores representing the standard error. Subjects were asked to rate each sensation on a 0 (no sensation) to 10 (max sensation) scale.

| During tACS |  |  | After tACS |  |  |
| --- | --- | --- | --- | --- | --- |
| Measure | Alpha | Beta | Measure | Alpha | Beta |
| Tingling | **2.3** | **3.4** | Tingling | **0.3** | **0.4** |
|  | *2.4* | *3.0* |  | *0.7* | *1.1* |
| Itching sensation | **2.3** | **2.9** | Itching sensation | **0.4** | **0.5** |
|  | *2.4* | *2.5* |  | *1.0* | *1.0* |
| Burning sensation | **1.0** | **1.1** | Burning sensation | **0.1** | **0.2** |
|  | *1.8* | *1.9* |  | *0.3* | *1.0* |
| Pain | **0.5** | **0.2** | Pain | **0.1** | **0.0** |
|  | *1.2* | *0.3* |  | *0.3* | *0.1* |
| Fatigue | **0.5** | **1.2** | Fatigue | **0.6** | **0.5** |
|  | *0.9* | *2.2* |  | *0.9* | *1.1* |
| Nervousness | **0.3** | **0.2** | Nervousness | **0.0** | **0.0** |
|  | *0.7* | *0.5* |  | *0.1* | *0.1* |
| Headache | **0.3** | **0.0** | Headache | **0.1** | **0.1** |
|  | *0.7* | *0.1* |  | *0.4* | *0.3* |
| Difficulty concentrating | **2.2** | **2.0** | Difficulty concentrating | **0.1** | **0.2** |
|  | *2.7* | *2.2* |  | *0.3* | *0.5* |
| Mood change | **0.3** | **0.1** | Mood change | **0.0** | **0.0** |
|  | *0.6* | *0.2* |  | *0.1* | *0.1* |
| Change in your vision/visual perception | **3.0** | **2.9** | Change in your vision/visual perception | **0.2** | **0.1** |
|  | *3.0* | *2.9* |  | *0.5* | *0.2* |
| Visual sensation | **3.6** | **5.5** | Visual sensation | **0.0** | **0.2** |
|  | *3.9* | *2.7* |  | *0.2* | *0.4* |

## Footnotes

1 Conducted using the *vartest2* function in Matlab Statistics Toolbox

